# PFKP regulates AXL-MET oncogenic and metabolic pathways in lung cancer

**DOI:** 10.1101/2024.03.03.583230

**Authors:** Yuze Sun, Huijie Zhao, Huijing Feng, Yue Liu, Yu Li, Sijie Chen, Zhiqing Zhou, Yuhui Du, Xiaofei Zeng, Huan Ren, Wenmei Su, Qi Mei, Guoan Chen

## Abstract

**Objective:** PFKP (Phosphofructokinase, Platelet Type isoform), as an essential metabolic enzyme, contributes to the high glycolysis rates seen in cancers, while its role in oncogenic pathways, especially from a non-metabolic aspect, is not fully understood.

**Methods:** Here we performed a comprehensive analysis of published RNA-seq, microarray data, and immunohistochemistry of tissue microarray to evaluate the significance of PFKP expression in non-small cell lung cancer (NSCLC). Functionally, we tested the cell proliferation, colony formation, invasion, and migration upon PFKP knockdown in lung cancer cells. Mechanistically, we performed RNA-seq, DIA-mass spectrum, western blot, and qPCR to probe the change of cell signaling pathways upon PFKP silencing. Co-immunoprecipitation and mass spectrum were used to uncover potential PFKP interacting proteins.

**Results:** We found that PFKP was highly expressed in NSCLC and was related to poor patient survival. Knockdown of PFKP significantly inhibited cell proliferation, colony formation, invasion, and migration of NSCLC cells. Mechanistically, we found that PFKP can directly bind with AXL and promote its phosphorylation at Y779, thus activating the AXL signaling pathway and promoting MET phosphorylation. In addition, several glycolysis, glutaminolysis, and TCA cycle proteins were downregulated following PFKP silencing.

**Conclusions:** These data demonstrate that PFKP, beyond its known role in glycolysis, also has a distinct non-metabolic function in affecting lung cancer progression by directly interacting with the AXL-MET axis, thus indicating a potential therapeutic target for lung cancer.

## 1. Introduction

Lung cancer is the leading cause of cancer-related deaths in both males and females, and almost one-quarter of all cancer deaths are due to lung cancer [1; 2]. Non-small cell lung cancer (NSCLC) accounts for more than 80% of all lung cancers. Despite improvement with new clinical management, the 5-year survival rate of lung cancer still remains only 19%[1]. Cancer often demonstrates distinct metabolic properties, with tumor cells preferring to perform anabolic metabolism in spite of its low energy conversion efficiency[3]. Several studies have examined how this anabolic preference contributes to cancer progression[4], and that many glycolysis related enzymes have also been reported to have non-metabolic activities that are essential for cellular function as well as cancer progression[5], suggesting hyperactivation of the glycolytic pathway in cancer has more than just metabolic consequences. However, the mechanism of how the hyperactivation of glycolytic pathway might contribute to cancer related signaling has not been fully understood.

PFKP (Phosphofructokinase, Platelet Type) is one of the three isoforms of PFK1, an enzyme that catalyzes the rate limiting step of glycolysis, and its high expression strongly promotes cell anabolic metabolism[6]. PFK1 is a tetramer composed of PFKP (platelet isoform), PFKM (muscle isoform) and PFKL (liver isoform), it can be either homo tetramer or hetero tetramer, the composition of the PFK1 tetramer varies depending on the tissue and cell types[7]. Of these three isoforms, PFKP has been identified to have distinct high expression in tumors[8; 9]. Recently, PFKP has been reported to be both a mediator of cancer cell metabolism and promote the development and progression of many types of cancer [10], including breast cancer[11; 12], oral cancer[13], glioblastoma[14; 15], kidney cancer[16] and lung cancer[17; 18]. Though the biological functions of PFKP in lung cancer have been elucidated, the mechanism of its relationship with specific oncogenic pathways is not fully understood.

Receptor tyrosine kinases (RTKs) play an important role in a variety of cellular processes including cell proliferation, migration, metabolism, and its dysregulation is tightly related to multiple types of cancers including NSCLC[19]. MET (MET proto-oncogene, Receptor Tyrosine Kinase) and AXL (AXL Receptor Tyrosine Kinase, Anexelekto) belong to RTK family of kinases and are important therapeutic targets for NSCLC[20; 21]. AXL has two phosphorylation sites necessary for its activation, Y702 and Y779. Previous studies have shown that Y702 mainly undergoes ligand-dependent autophosphorylation while Y779 phosphorylation involves interaction of AXL with other kinases[22]. The ligand independent Y779 phosphorylation of AXL can be triggered by other types of RTKs, including EGFR (Epidermal Growth Factor Receptor)[23], ErBb2[24], ErBb3[25] and MET[26; 27], while the activated AXL can also promote the activation of the interacting RTK. This heterodimerization-dependent activation mechanism of RTKs is widespread in cancers and is important for both cancer development and progression.

In this study, we demonstrated that PFKP could bind with AXL and promote its phosphorylation at Y779, which may further activate MET via the heterodimerization of MET and AXL. In addition, several glycolysis, glutaminolysis and TCA cycle proteins were decreased after PFKP knockdown. These results indicate that PFKP may play a critical role in lung cancer progression not only through the metabolic pathway, but also via an oncogenic mechanism. This could potentially provide a new therapeutic strategy for lung cancer therapy.

## 2. Materials and methods

### 2.1. Cell lines and cell culture

A549, H1299 and H838 cell lines were obtained from ATCC (Supplementary **Table S1**). All cell lines were maintained in RPMI 1640 (Gibco) supplemented with 10% FBS (Gibco) and 1% antibiotic agents and cultured at 37_ in a 5% CO2 cell culture incubator. For cell passage, after discarding the medium, plates were washed with PBS (Gibco, pH 7.4), digested using 0.05% trypsin-EDTA (Gibco) for 5 min at 37_, before stopping the digestion by adding a double volume of medium. The cells were then centrifuged at 1000 RCF for 3 min, the supernatant discarded, and the cells then resuspended using medium before addition to new plates with complete medium for further culture.

### 2.2. Published microarray and RNA-Seq data collections

Four groups of microarray data were downloaded from NCBI Gene Expression Omnibus (Hou:GSE19188[28], Shedden: GSE68465[29], Landi: GSE10072[30]), TCGA and GTEx data were downloaded from GEPIA[31]. RNA-seq datasets were downloaded from three publications including Seo (87 AD)[32], Collisson (312 AD)[33] and Dhanasekaran (67 AD)[34]. Expression levels of transcripts were represented as reads per kilobase per million mapped reads (RPKM)[35].

### 2.3. siRNA mediated knockdown

siRNAs were obtained from Genepharma, the sequences of siRNA are listed in Supplementary **Table S2**. Cells were plated as desired concentration 8 hrs before transfection. siRNA was dissolved in ddH_2_O to a concentration of 1μM before use. RNA iMAX (Invitrogen) and siRNA were dissolved in OptiMEM medium (Gibco) according to the manufacturer’s instructions. The working concentration of siRNA is 10nM. The knockdown efficiency was tested by both Western blot and qPCR.

### 2.4. Lentiviral transfection

PFKP expression lentiviral was purchased from HANBIO, the vector is LV011-PHBLV-CMV-MCS-3FLAG-EF1-T2A-Zsgreen-Puro. PFKP gene was labeled with 3x Flag tag, and the vector expresses Zsgreen as well. Transfection was performed in accordance with the manufacturer’s instructions. H1299 cells transfected with Flag-PFKP overexpression or empty vector (negative control) were created as a stable cell line.

### 2.5. Cell proliferation assay and colony formation assay

For cell proliferation assay, cells were plated in 96 well plate at the desired concentrations, and siRNA knock downs were performed 8hrs after plating, either using PFKP targeted siRNA or non-target siRNA. Five repeats were done for each group. Cell viability was tested after 96hrs of transfection using MTS kit (Promega) according to the manufacturer’s instructions.

For colony formation assay, cells transfected with either PFKP targeted siRNA or non-targeted siRNA were seeded in 6-well plates at a concentration of 500 cells per well. Cells were cultured at 37_ for ten days and medium was then discarded, fixed in 100% methanol, stained with 0.1% crystal violet (Solarbio), and the number of colonies counted in each well.

### 2.6. Cell invasion and migration assay

For invasion and migration assays, cells were treated with siRNA for 48 hrs and transferred to chambers on 24 well plates at 10^5^/0.5ml per well. The chambers were plated with (invasion assay) or without (migration assay) matrigel, 10% FBS was added to the lower chamber as a chemoattractant, and the medium above was FBS free. Cells were incubated at 37_ for 24 hrs, and the chambers were washed and fixed, and stained with crystal violet for observation and photographed under the microscope. All the groups were performed in triplicate.

For the wound healing assay, cells were plated in 6 well plates at 5 *10^8^ per well, treated with siRNA, and cells became confluent after 48 hrs. Following scratching the plate with pipette tips, the plates were washed with PBS and FBS free medium added. The pictures of the plates before and 24 hrs later were recorded. All the groups were performed in triplicate.

### 2.7. RNA extraction and quantitative real-time RT-PCR

Forty-eight hrs after siRNA mediated knock down, cells were digested, collected and RNA isolated using Trizol (Ambion), following the manufacturer’s instructions. The RNA was dissolved in DEPC-treated H2O (65 _ for 10-15 min) and the concentration determined using nanodrop (Thermo). The cDNA synthesis and gDNA erase protocol were performed using the PrimeScript RT reagent Kit with gDNA Eraser (Takara). Quantitative real-time RT-PCR (qRT-PCR) was performed using TB Green Premix Ex Taq ™ II (Tli RNaseH Plus) (Takara) with qTOWER3 (Jena). Each sample was analyzed in triplicate, and the housekeeping gene beta-actin was used as a loading control. The sequences of primers are listed in Supplementary **Table S3.**

### 2.8. RNA-seq

For RNA-seq, extracted RNA was sent to Genedenovo Biotechnology Co., Ltd (Guangzhou, China) for RNA-seq analysis. After enrichment of poly A+ mRNA using magnetic beads with oligo (DT), the isolated mRNA was fragmented by ultrasound. The first cDNA strand was synthesized by reverse transcriptase (PCR) and the second strand was synthesized by reverse transcription polymerase chain (PCR). The purified double stranded cDNA was treated with terminal repair, A-tail and sequencing adaptor. The 200 bp cDNA was screened by AMPure XP beads and PCR amplification was performed. The PCR product was purified by AMPure XP beads, and finally the library was obtained and an Agilent 2100 Bioanalyzer was used to detect RNA integrity.

FASTP[36] was used to do quality control for raw reads, the steps of reads filtering are as follows: 1) Remove reads with adapter. 2) Removal of reads containing more than 10% N. 3) Remove all a-Base reads. 4) Remove low quality reads (base with mass value Q ≤ 20 accounts for more than 50% of the whole read). Subsequently, bowtie 2[37], a short reads alignment tool, was used to align reads to the ribosomal database. HISAT2[38] was used to carry out comparative analysis based on reference genome. According to the comparison results of hisat2, STRINGTIE[39] was used to reconstruct the transcripts and RSEM[40] to calculate the expression levels of all genes in each sample. Analysis of differences between groups were performed by DESeq2[41].

### 2.9. Protein extraction and immunoblot analysis

The cells were lysed in RIPA buffer (VETEC) supplemented with protease inhibitor PMSF (Cell Signaling Tree). Cell lysate was centrifuged, protein concentrations were determined using Pierce BCA Protein assay kit (Thermo), adding 4x SDS loading buffer, and then denatured at 100_ for 10 min. The proteins were electrophoresed by pre-made SDS page (GenScript) and transferred to PVDF membranes (Roche). The membranes were blocked in 5% non-fat milk (Difco) and then incubated with primary antibodies overnight at 4_. The next day, membranes were incubated with secondary antibody (Cell Signaling Tree) for 1h at room temperature, after washing with TBST (Boster), HRP substrate (Immobilon) was added prior to performing exposure. Antibodies used in the study are listed in Supplementary **Table S4**.

### 2.10. Co-immunoprecipitation and mass spectrum analysis

The H1299 cell line with Flag labeled PFKP overexpression or empty vector (negative control) were used to prepare lysates. IP lysis buffer (Thermo), anti-flag magnetic beads (Bimake) were added to the cell lysate, and subsequent procedures were done according to manufacturer’s instructions. For AXL pull down, AXL (C89E7) (Cell Signaling Tree) antibody was added to the cell lysate, incubated with shaking overnight, and magnetic beads (Invitrogen) were added to the lysate, and subsequent steps done according to manufacturer’s instructions. Beads were subsequently used for Western blot analysis and spectrum analysis.

For spectrum analysis, beads after washing were sent to Wininnovate Bio for 60 min mass spectrum analysis. Proteins having 10-fold higher peak areas in FLAG-PFKP group compared to the NC group were selected as potential significant interactors of PFKP.

### 2.11. DIA-Mass Spectrometry

Proteins were identified and quantified using a data independent acquisition (DIA) based Mass Spectrometry (MS) method. A spectral library was created by analyzing six times with data dependent acquisition method (DDA). Sample preparation involved the process of protein denaturation, reduction, alkylation, tryptic digestion and peptide cleanup. Analytical separation was conducted on an Ultimate 3000 HPLC system (ThermoScientific). Peptides were analyzed on a Fusion Orbitrap mass spectrometer (ThermoScientific) equipped with an Easy-nLC 1200 (ThermoScientific). Raw Data were processed and analyzed by Spectronaut X (Biognosys AG, Switzerland) with default settings to generate an initial target list. Qvalue (FDR) cut off on precursor and protein level was applied at 1%. DIA-MS was performed by Genedenovo Biotechnology Co., Ltd (Guangzhou, China).

### 2.12. Immunohistochemistry of tissue microarray

Lung cancer tissue microarray (TMA) (HLugA180Su08 and HLugA150CS03) was provided by Outdo Biotech Company. The TMA used in this study was approved by the Ethics Committee of Shanghai Outdo Biotech Company. After the 4 μM slices on slides were dewaxed in xylene and rehydrated in an ethanol series to water, antigen retrieval was performed with sodium citrate. One sachet of sodium citrate (Solarbio) was dissolved in 2L PBS according to the instructions and heated to boiling in a microwave oven. Then the slides were added with stopping for 1 min for every 5 min of heating at 60% heat, repeated 3 times, and cooled naturally to room temperature. Endogenous peroxidase was blocked with hydrogen peroxide, and 10% BSA + 10% goat serum was used to block the non-specific binding to other antigens. The slides were incubated with PFKP antibody (Cell Signaling Technology, 1:50 dilution) for 1hr at 37℃, followed by antibody detection with a goat anti-rabbit DAB detection kit (MaxVision).

### 2.13. Statistical analysis

All data analysis was performed using GraphPad Prism 8 (GraphPad software) and Excel. Kaplan-Meier survival curves were done in the GraphPad Prism 8. Heatmaps were done in cluster 3.0 software and viewed in Tree View software. Data from proliferation assays and colony formation assays were analyzed using unpaired t-test, with a p value <0.05 was considered statistically significant. Metascape[42] and DAVID[43] were used for GO, KEGG and Reactome enrichment analysis, with the p value cut off set as 0.01.

## 3. Results

### 3.1 PFKP is a glycolytic gene highly expressed in cancer tissues and its high expression is associated with poor patient survival

Cancer cells show a preferential dependence on glycolysis, even though it is less efficient than oxidative phosphorylation in terms of ATP production. This oxygen-independent manner, called the Warburg effect, is essential for cancer origination and progression[3]. This distinct glycolysis pathway in cancer has been proposed as a target for cancer therapy[44]. However, the specific mechanism underlying the high activation of the glycolysis pathway favoring cancer progression, and particularly those involving non-metabolism-related aspects, have not been elucidated. In order to find the potential links between high-level glycolysis and cancer progression, we analyzed the expression of 56 glycolytic genes in lung cancer samples from TCGA [45] lung adenocarcinomas (LUADs) and GTEx [46] normal lung tissues data to identify glycolytic genes that differ significantly between normal lung and lung tumor tissues. (**Figure 1A** and **Table S5**). Next, by combining data from Shedden et al (442 LUADs [29]), we performed gene screening using the following three criteria: genes with greater than 2-fold increase in expression in tumors as compared to normal tissues, high expression of the genes associated with unfavorable for patient survival with p value <0.05, and genes highly expressed in lymph node metastasizing tumors as compared with non-lymph node metastasizing tumors with p value < 0.05. We found that PFKP met all the three criteria very well. PFKP mRNA shows higher expression in LUAD (**Figure 1A**), higher expression in tumors with lymph node metastatic status (**Figure 1B**) and higher PFKP level was unfavorable for patient survival (**Figure 1C**). We next examined PFKP expression in other datasets, and found PFKP expression was also significantly higher in tumors in data from Hou et al [28](**Figure 1D**)[28] and Landi et al [30] (**Figure 1E**). In Shedden data, PFKP mRNA was higher in tumors with Stage 2 and 3 as compared to Stage 1(**Figure 1F**), and higher in tumors with moderate and poor differentiation as compared to well differentiation (**Figure 1G**). In addition, we further verified using TCGA data and observed that PFKP mRNA expression was higher in LUAD and LUSC (compared with matched normal lung tissues) (**Figure 1H**) and higher expression was associated with poor survival in LUAD (**Figure 1I**) and other multiple types of tumors in TCGA data (**Figures S1A-S1G**).

**Figure 1.**
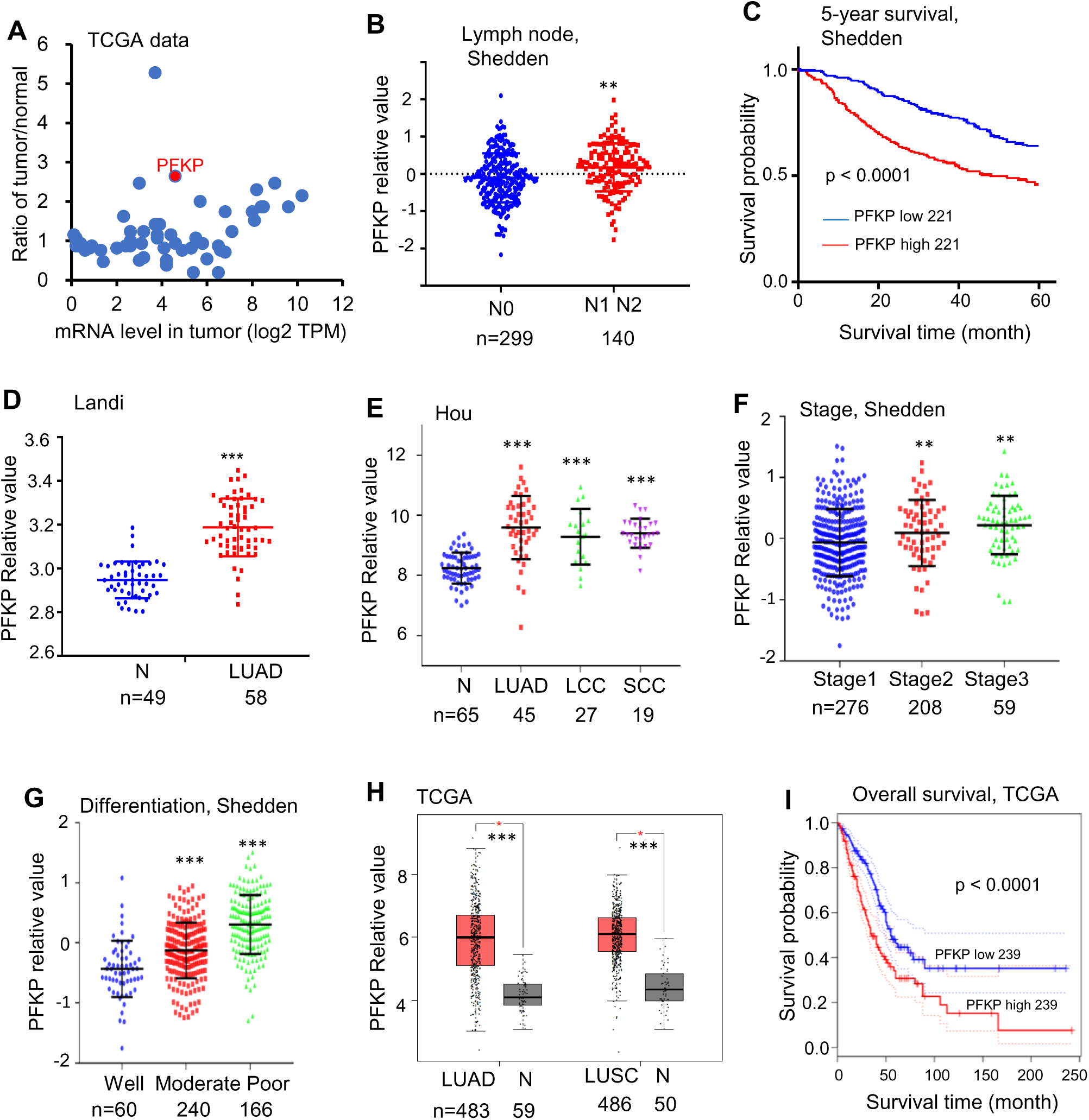
PFKP shows high expression in lung cancer and correlates to poor patient prognosis. **A**, scatter plot of mRNA expression for 56 glycolytic genes in lung adenocarcinomas (LUADs) and fold changes of tumor/normal (data from TCGA LUAD and GTEx normal); **B**, PFKP mRNA expression in tumors with lymph node metastasis status (data from Shedden et al); **C**, Kaplan-Meier survival curves with a log-rank test of PFKP in Shedden data (442 LUADs); **D, E**, PFKP mRNA expression levels in lung tumor (T) and normal (N) tissue samples in data form Landi et al and Hou et al. LCC (large cell) and SCC (squamous cell), ***: p<0.001; **F, G**, PFKP mRNA expression in tumors with stage and differentiation (data from Shedden et al with 442 LUADs samples), **: p<0.01, ***: p<0.001; H, PFKP mRNA expression levels in LUAD, LUSC and with matched normal tissue samples in data from TCGA (GEPIA web: http://gepia.cancer-pku.cn); **I,** Kaplan-Meier survival curves with log-rank test of PFKP in TCGA (GEPIA web) data.

PFKP protein expression data from primary lung tumors measured by mass spectrometry was available on the UALCAN website[47; 48]. It was higher in lung adenocarcinoma as compared to normal lung tissue, as well as higher in poorly differentiated tumors (**Figures S1H** and **S1I**). These data demonstrated that PFKP, as a glycolytic gene, may have an essential oncogenic role in cancers.

### 3.2. PFKP protein expression is higher in human lung cancer specimens and mRNA correlated genes are involved in cell cycle, glycolysis and cancer related pathways

To determine if PFKP is important in human lung cancer progression, we performed PFKP protein expression and KEGG pathway analysis based on human lung specimens. We first performed immunohistochemical (IHC) staining for PFKP protein using a lung tissue microarray containing 98 lung adenocarcinoma samples (**Figure S2A**). The results showed that PFKP protein was highly expressed in lung adenocarcinomas (**Figures 2A-G**), and higher expression of the PFKP protein was related to poor patient survival in lung cancer (**Figure 5H**). PFKP protein staining was mainly present in the cytoplasm which was consistent with Western blotting in **Figure S2B**.

**Figure 2.**
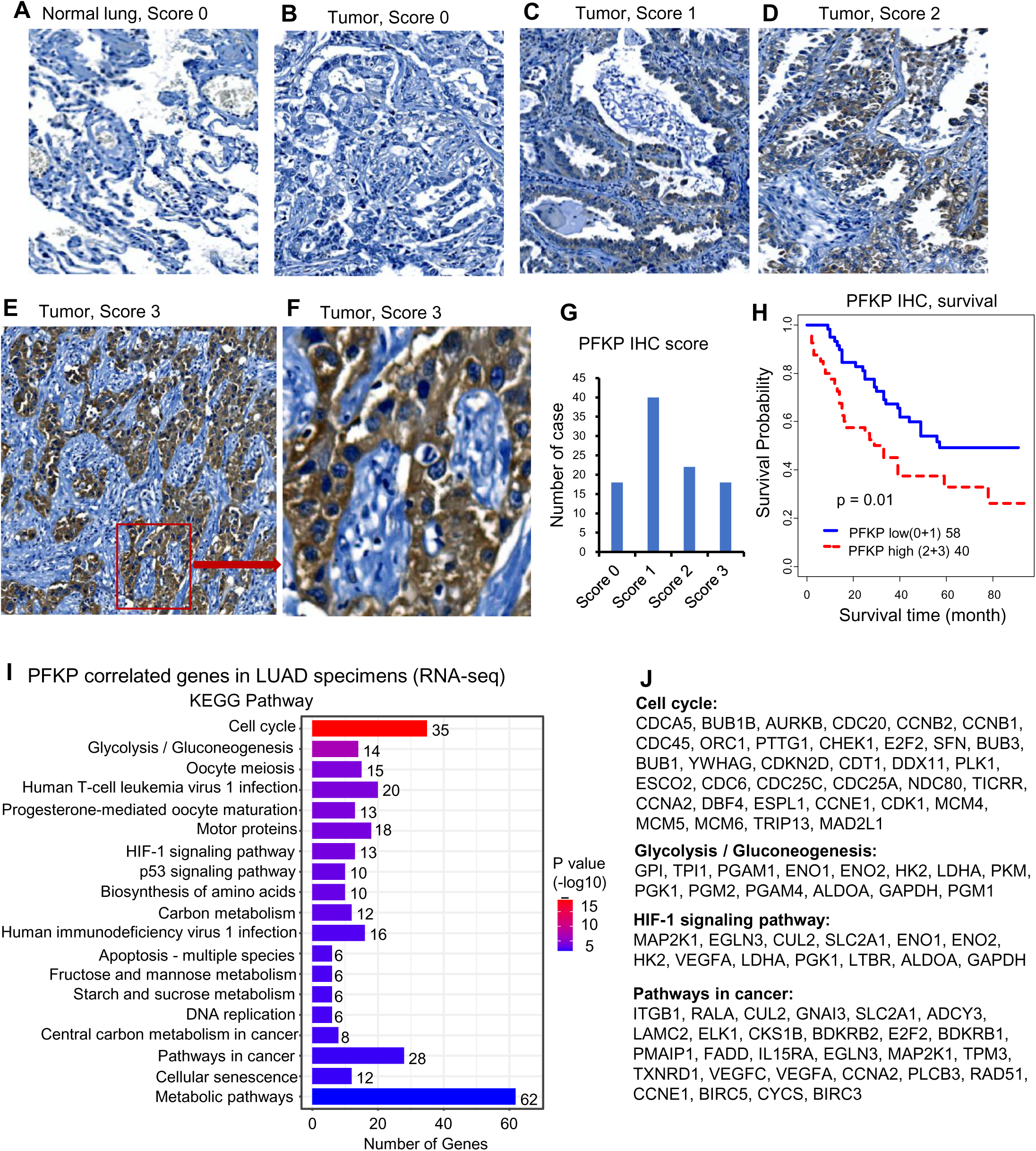
PFKP protein expression is higher in human lung cancer specimens and mRNA correlated genes are involved in cell cycle, glycolysis and cancer related pathways. **A-F**, representative images of IHC from lung TMA with score 0-3; **G**, Summary of IHC score of the TMA; **H**, Kaplan-Meier survival curves with log-rank test of PFKP protein expression from IHC of the TMA (score 0+1 vs. 2+3); **I**, KEGG pathway analysis based on PFKP correlated genes from LUAD specimens; **J**, genes in several important KEGG pathways.

Next, we performed the Pearson correlation analysis between PFKP and other genes (mRNA) using three LUAD datasets including Seo (87 AD)[32], Collisson (312 AD)[33] and Dhanasekaran (67 AD)[34]. We selected the correlated genes based on the average r value of these 3 datasets. There were 564 genes significantly positively correlated to PFKP in LUAD (Pearson correlation r >=0.25, n=466, p <0.001). We performed the KEGG pathway analysis using DAVID website[43] using these 564 genes. The results showing that cell cycle and glycolysis/gluconeogenesis pathways were on the top list. HIF-1 signaling, P53 signaling, DNA replication, pathways in cancer and metabolic pathways were also involved in PFKP related genes (**Figure 2I**). The genes involved in cell cycle, glycolysis/gluconeogenesis, HIF-1 and pathways in cancer were listed in **Figure 2J**. These results indicate PFKP may play an important role in the clinical behavior of human lung cancer.

### 3.3. PFKP plays an oncogenic role in lung cancer progression *in vitro*

To explore the biological function of PFKP on lung cancer cells *in vitro*, we knocked down PFKP using siRNAs and verified its knockdown efficiency by Western blot (**Figure 3A**). We found that the cell proliferation and colony formation were significantly inhibited upon PFKP knockdown (**Figures 3B-F**) as previously reported[17]. Knocking down PFKP in NSCLC has been reported to decrease cell invasion and migration of H1299 and A549[18]. We performed a transwell assay and wound healing assay using the H838 cell line, and found that cell invasion through Matrigel-coated membranes was significantly decreased upon PFKP knockdown as compared with the cells treated with nontarget control siRNA. Cell migration was also inhibited after knockdown of PFKP (**Figures 3G** and **3H**). In line with transwell assay results, the wound healing assay showed that cells having PFKP silenced have a lower speed of healing compared with no target siRNAs transfected group (**Figures S3A** and **S3B**). In contrast, the rate of wound healing in H1299 cells was accelerated after PFKP overexpression (**Figures S3C** and **S3D**). These data suggest that PFKP contributes to lung cancer progression and metastasis through increases in cell proliferation, invasion and migration.

**Figure 3.**
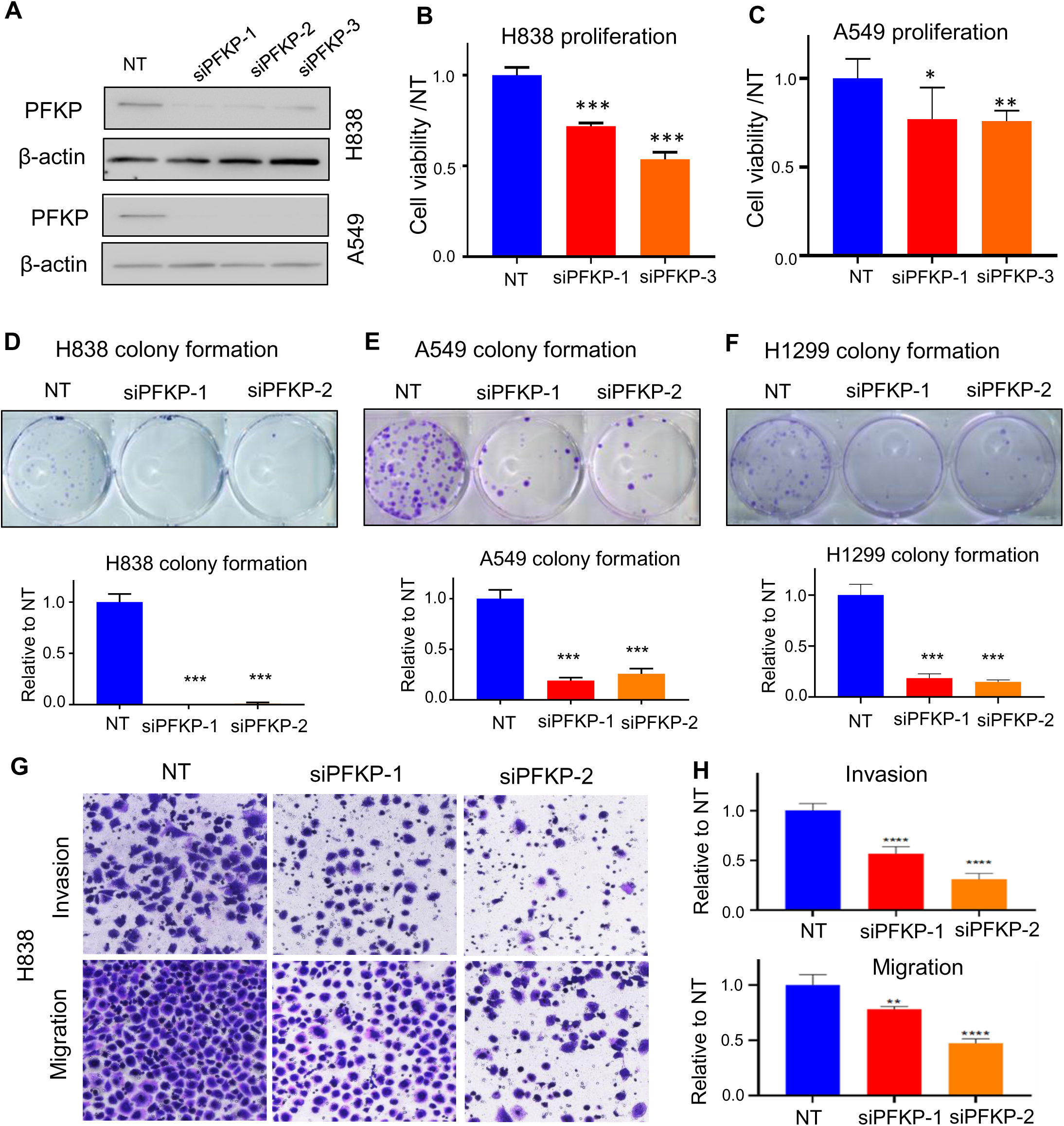
PFKP affects tumor cell proliferation, colony formation, migration and invasion. **A**, siRNA efficiently was able to knockdown PFKP in A549 and H838 cell lines (72h) measured by Western blot; **B, C**, knockdown of PFKP significantly inhibited cell proliferation in A549 and H838, * p<0.05, ** p<0.01, *** p<0.001; **D-F**, knockdown of PFKP by siRNA lead to decrease in colony formation in H838, A549 and H1299, low panel is the relative quantified value of up panel, *** p<0.001; **G**, **H**, siRNA mediated PFKP knockdown leads to decreased cell migration and invasion in H838, relative value on the right (**H**), *** p<0.001.

### 3.4. AXL directly binding to both PFKP and MET

PFKP has been reported to directly interact with EGFR[49], which is also a receptor tyrosine kinase. In order to define potential PFKP interacting proteins, we performed co-immunoprecipitation (Co-IP) and mass spectrum analyses. In H1299 cells transfected with Flag-labeled PFKP, we performed pull down of PFKP using anti Flag magnetic beads, and the interacting proteins were then measured by mass spectrometry. We found that there are 157 proteins having 1.5-fold change (area over-flag-PFKP/vector) with p value >20 (-log10) (Supplementary **Table S6**). To our surprise, EGFR protein was not presented in the list of PFKP interactors, but another receptor tyrosine kinase, tyrosine-protein kinase receptor UFO (AXL) emerged (**Figure 4A**, Supplementary **Table S6**). We also found that glucose transport GLUT1 and lactate transport MCT1 may potentially bind directly to PFKP (Supplementary **Figure S4A-C,** Supplementary **Table S6**). In addition, several solute carrier family members and several ras-related proteins were uncovered with this PFKP Co-IP MS assay (Supplementary **Table S6**). AXL, a member of the TAM (Tyro3, Axl, Mer) family, has been reported to have oncogenic functions in multiple types of cancers and is involved in many signal transduction cascades in response to its ligand growth arrest specific 6 (GAS 6)[50]. GAS 6 activation is related to EGFR inhibitor resistance in lung cancer[51] and a therapeutic target for clinical treatment[52]. We further confirmed the interaction between PFKP and AXL using Co-IP with Flag antibody and Western blot assays in H1299 cells transfected with Flag-PFKP. As shown in **Figure 4B**, PFKP binds to AXL but not MET. It was reported that AXL and MET can interact with each other [26; 27], next, we performed Co-IP pull down using an AXL antibody, and in line with our Flag pull down result, it shows that AXL interacts with both PFKP and MET directly in H1299 cells transfected with Flag-PFKP (**Figure 4C**), or in H1299 cells without Flag-PFKP (**Figure 4D**). These results suggest that AXL binds with AXL and MET directly in NSCLC.

**Figure 4.**
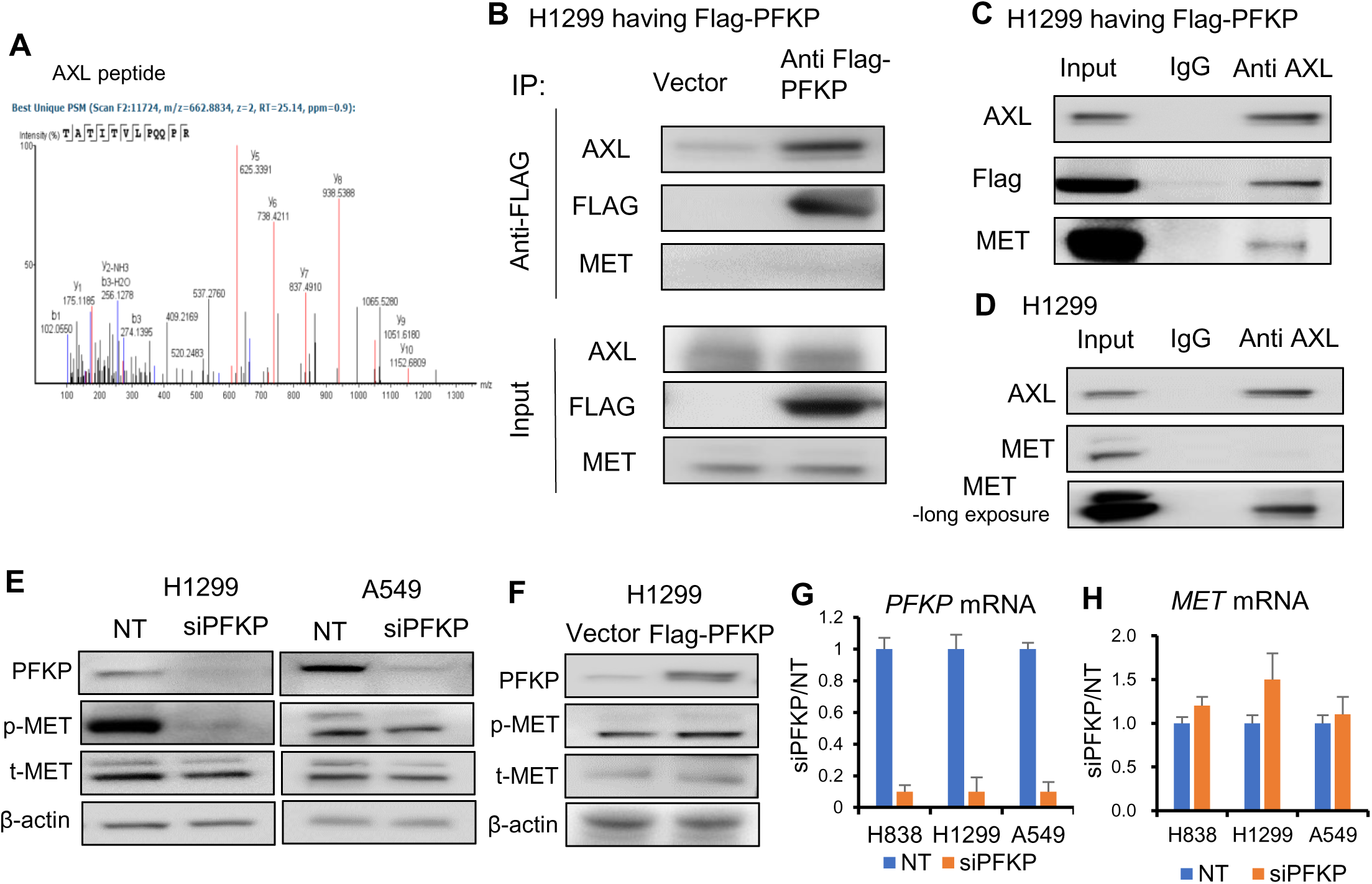
AXL directly binding to both PFKP and MET. **A**, PFKP Co-immunoprecipitation (Co-IP) followed by mass spectrometry indicate that the peptide of AXL was pulled down by PFKP; **B**, Western blot after pull down using anti-flag antibody Co-IP verified that PFKP binds to AXL directly but not MET; **C**, AXL Co-IP pull down using anti-AXL antibody shows that AXL interacts with PFKP and MET in H1299 cells with Flag-PFKP; **D**, AXL Co-IP pull down using anti-AXL antibody shows that AXL binds to MET directly in H1299 cells; **E**, PFKP silencing by siRNA leads to p-MET protein decrease in H1299 and A549 cell lines; **F**, overexpression of PFKP increases p-MET protein in H1299 cells; **G,** siRNA efficiently knockdown PFKP in lung cancer cell lines (48h) measured by qRT-PCR; **H**, *MET* mRNA was not changed after PFKP knockdown measured by qRT-PCR.

By Western blot analysis, we found that MET Tyr 1234/5 phosphorylation was significantly suppressed in H1299 and A549 cell lines (**Figure 4E**). Further, we stably overexpressed PFKP using lentiviral transfection in H1299, which had relatively low expression of PFKP (**Figure S4D**, data from CCLE[53]), and found that MET phosphorylation level was increased upon overexpression of PFKP (**Figure 4F**). MET is a potential therapeutic target in NSCLC, and can promote lung cancer progression through multiple mechanisms including increased cancer cell survival, growth, and invasiveness[54]. MET mutations are prevalent in NSCLC patient tumors[21]. It was reported that MET phosphorylation at Tyr 1234/5 is essential for MET pathway activation[55]. While, MET mRNA level remained unchanged upon PFKP knockdown as measured by RT-PCR (**Figures 4G** and **4H**), suggested that PFKP regulate MET may through MET phosphorylation rather than via transcriptional regulation.

We also tested several other oncogenic proteins such as STAT3, mTOR, NF-kB, NOTCH1, AKT, KRAS and RAF and no changing were found after PFKP silencing (**Figures S4E**).

### 3.5. PFKP promotes MET phosphorylation via enhancing AXL 779Y phosphorylation

We found that PFKP directly interacts with AXL and promotes MET phosphorylation. It was previously reported that AXL and MET interact with each other, and this interaction can promote phosphorylation of both receptors[26; 27], referred to as the receptor tyrosine kinase hetero interaction. We thus postulated that PFKP promotes MET phosphorylation through an AXL dependent mechanism rather than direct interaction. We performed Co-IP using anti-AXL pull down and found that, as previously reported, AXL binds to MET directly (**Figure 4C** and **4D**). We also found that PFKP protein expression was only present in the cytoplasm (**Figure S2B**).

AXL is known to have mainly two phosphorylation sites for its activation, which are Y702 and Y779. Phosphorylation at Y702 is thought to be related to ligands for AXL, GAS6, whereas the Y779 phosphorylation was found to be ligand independent and can heterodimerize with many other kinases, including MET[27], EGFR[56], ErBb3[25] and HER2[24]. To test PFKP’s effect on AXL activation, we performed Western blot analyses and found that AXL Y779 phosphorylation was inhibited upon PFKP knockdown and increased following PFKP overexpression. By contrast, Y702 was not affected (**Figure 5A**). This implies that AXL phosphorylation at Y779 is related to MET phosphorylation. There was no significant change in AXL total protein or mRNA after PFKP knockdown (**Figure 5A** and **5B**).

**Figure 5.**
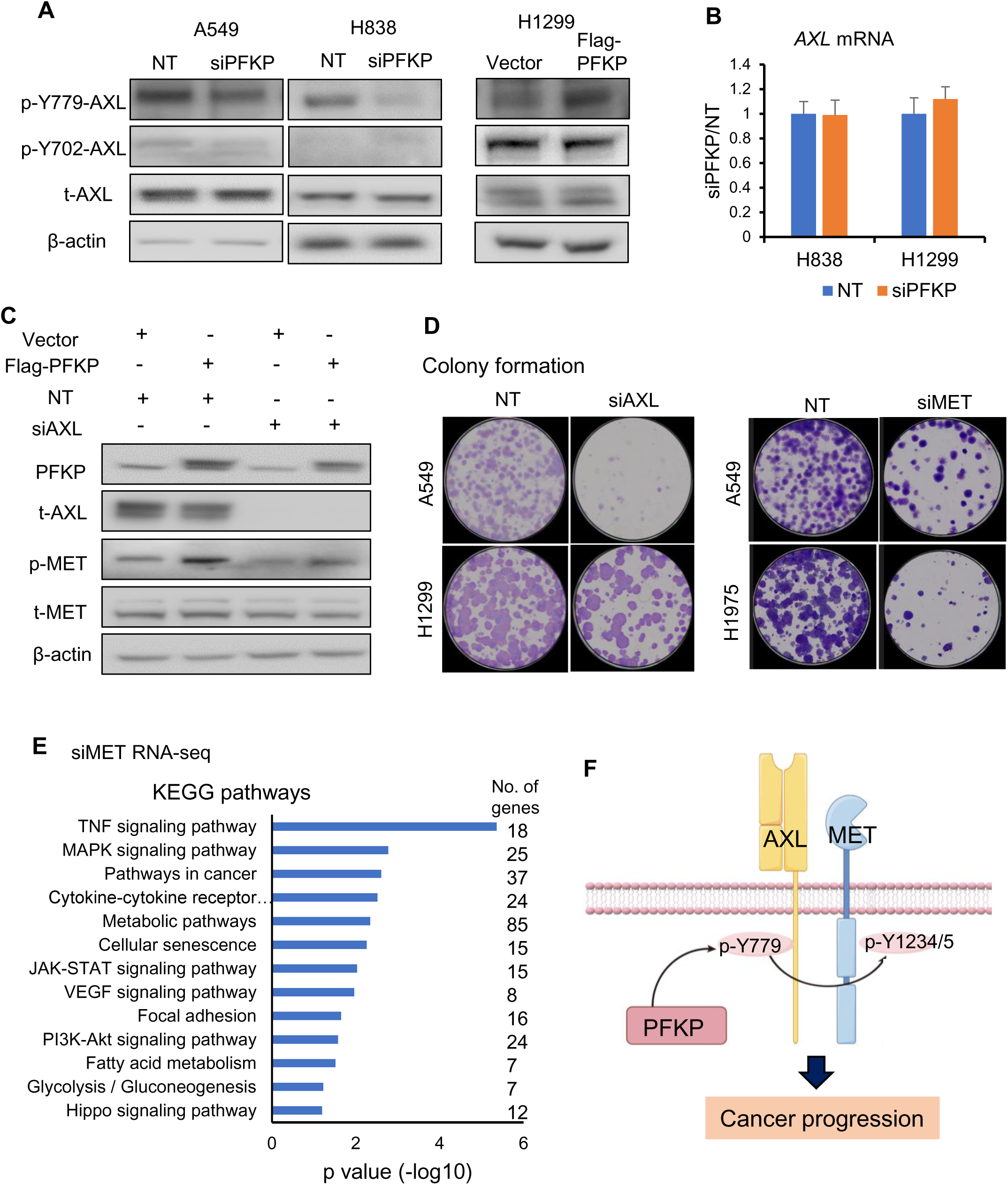
PFKP promotes MET phosphorylation via enhancing AXL 779Y phosphorylation. **A**, PFKP knockdown leads to AXL p-779 phosphorylated protein being decreased in A549 and H838 cells. Overexpression of PFKP (Flag-PFKP) leads to AXL p-779 phosphorylated protein increase while total AXL and p-702 proteins remain unchanged in H1299 cells; **B**, AXL mRNA was not changed upon PFKP knockdown; **C**, overexpression of PFKP leads to increase of phosphor-MET (p-MET), which is rescued by AXL knockdown; **D**, knockdown of AXL and MET by siRNAs lead to decrease in colony formation in lung cancer cells; **E**, KEGG analysis of 755 down regulated genes after MET knockdown as measured by RNA-seq in H1299, H1975 and PC-9 cell lines, genes changed (siMET/NT <0.65) in 2/3 cell lines; **F**, model of PFKP promotes MET phosphorylation by directly interacts with AXL and promotes its phosphorylation at Y779, and the activation of AXL and MET signaling pathway promote cancer progression.

To explore the relationship between PFKP’s regulation of AXL and MET, we performed rescue experiments. In H1299 cell line, phosphorylated MET increased upon PFKP overexpression, which was significantly rescued by siRNA mediated AXL knockdown (**Figure 5C**). Next, we performed colony formation assay to test the oncogenic role of AXL and MET, and found that the colony formation was decreased upon AXL or MET knockdown by siRNAs (**Figure 5D**). We also performed the RNA-seq and KEGG pathway analysis upon MET silencing by siRNAs in H1299, H1975 and PC-9 lung cancer cell lines. There were 755 genes decreased after MET knockdown. KEGG analysis of these genes indicated that TNF signaling pathway, MAPK signaling pathway, pathways in cancer and metabolic pathways were on the top list (**Figure 5E**) suggested that MET has roles in both oncogenic and metabolic processes. Taken together, these results demonstrated that PFKP promotes MET phosphorylation through direct binding to and enhancing AXL Y779 phosphorylation (**Figure 5F**). The mechanism of how PFKP regulates AXL Y779 phosphorylation is unknown and need further investigation.

### 3.6 Metabolism and cancer related signaling pathways are regulated by PFKP uncovered by RNA-seq and DIA-MS analysis in lung cancer cell lines

To further understand the potentially significant role of PFKP in lung cancer progression, we performed proteomics and genomic analysis using DIA-MS technology and RNA sequencing following PFKP knockdown with siRNAs in H838, H1299 and A549 lung cancer cell lines.

In the RNA-seq analysis, we chose genes with more than 1.5-fold changes as differential genes upon PFKP knockdown and selected overlapping genes in both H838 and H1299 cell lines. We obtained 510 down-regulated genes and 307 up-regulated genes upon silencing of PFKP (**Figures S6A-C**). Subsequently, we performed pathway enrichment analysis using the DAVID website[43] using these changed genes, and found that these 817 differential genes were mainly enriched in important metabolic related biologic processes and biological process (**Figure S6D**). In the proteomics analysis using DIA-MS technology upon PFKP knockdown, we found that there are 569 down-regulated proteins, and 558 up-regulated proteins after PFKP silencing in H1299, H838 and A549 cell lines. (Criteria: proteins changed (siPFKP/NT) in 2/3 cell lines <0.65, or up >1.5). KEGG pathway and GO BP analysis of these 1127 changed proteins revealed that these proteins were enriched in several cancer-related signaling pathways, organelle organization, catabolic processes and cell cycle pathways (**Figure 6A, B**). Importantly, we found that several glycolytic proteins such as GLUT1 (SLC2A1, Solute Carrier Family 2 Member 1), PGK1 (Phosphoglycerate Kinase 1), ENO2 (Enolase 2), PFKFB2/4 (6-Phosphofructo-2-Kinase/Fructose-2,6-Biphosphatase), were down-regulated after PFKP knockdown (**Figures 6C**). Meanwhile, several proteins involved in glutaminolysis and TCA cycle such as IDH3G (Isocitrate Dehydrogenase (NAD(+)) 3 Non-Catalytic Subunit Gamma), PDK2 (Pyruvate Dehydrogenase Kinase 2), LAT1 (SLC7A5, Solute Carrier Family 7 Member 5) and GLS (Glytaminase) were also decreased upon PFKP silencing (**Figures 6D**). When we looked at the mRNA levels of these genes from RNA-seq analysis, we found that the mRNAs of SLC2A1/GLUT1, PGK1, ENO2, IDH3G and SLC7A5/LAT1 were also decreased upon PFKP knockdown (**Figures 6E, F**). Both mRNA and protein of PFKM and PFKL were not affected by PFKP knockdown (**Figures S6E, F**).

**Figure 6.**
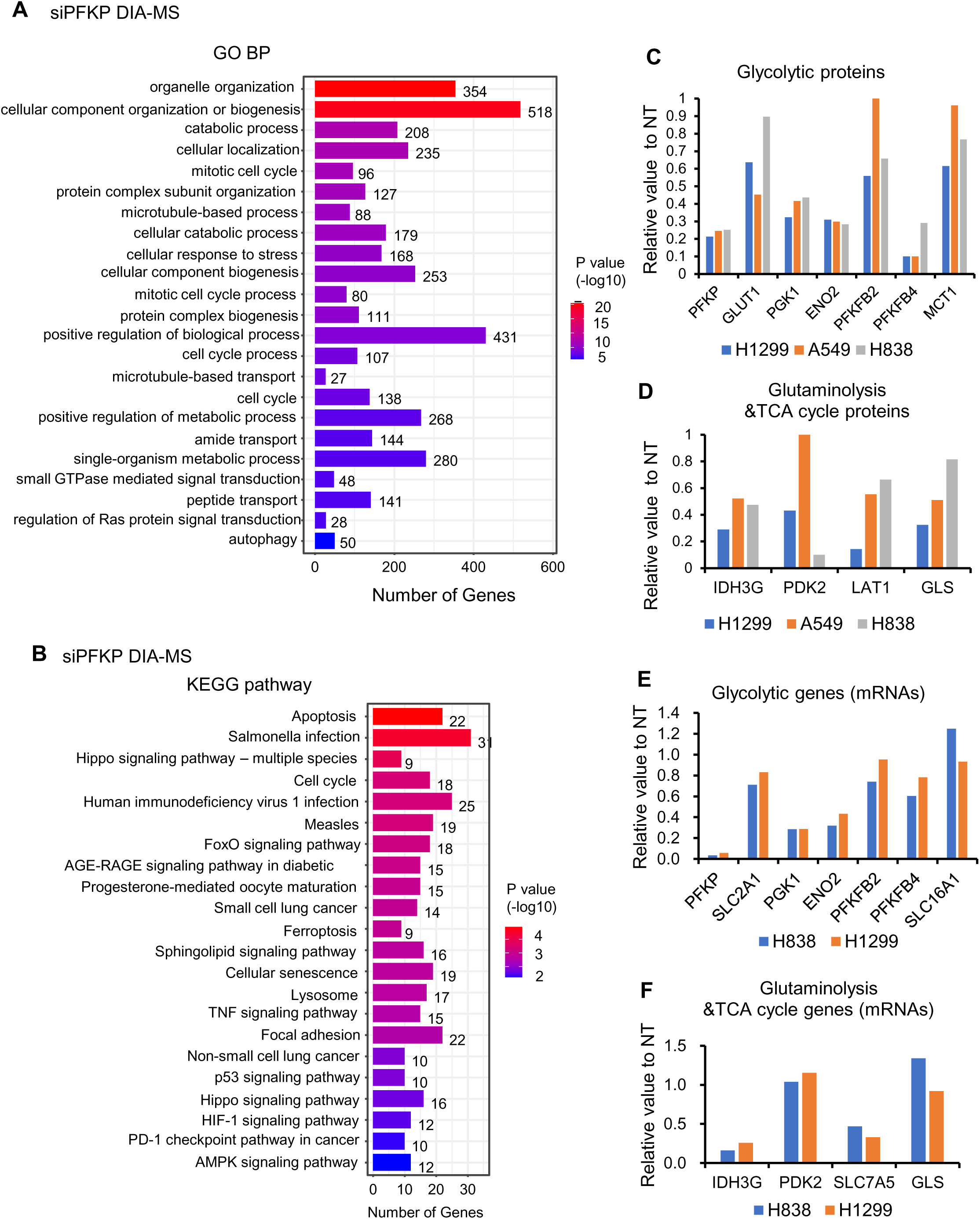
Metabolism and cancer related signaling pathways are regulated by PFKP uncovered by RNA-seq and DIA-MS analysis in lung cancer cell lines. GO BP (**A**) and KEGG (**B**) analysis of down and upregulated proteins after PFKP knockdown as measured by DIA-MS. H1299, H838, A549 proteins changed (siPFKP/NT) in 2/3 cell lines, down <0.65, 569 proteins, up >1.5, 558 proteins. **C** and **D,** several glycolytic proteins and glutaminolysis and TCA cycle proteins were decreased upon PFKP knockdown measured by DIA-MS in lung cancer cell lines; **E** and **F,** several glycolytic mRNAs glutaminolysis and TCA cycle mRNAs were decreased upon PFKP knockdown measured by RNA-seq in lung cancer cell lines.

Taken together, by combining DIA-MS, Co-IP and RNA-seq analysis, we found that several genes related to glycolysis, glutaminolysis and TCA cycle were regulated by PFKP (**Figure 7A**). Importantly, three major transporters, GLUT1 for glucose, LAT1 for glutamine and MCT1 for lactate were also regulated by PFKP through either directly binding and/or transcriptional regulation (**Figure 7A**).

**Figure 7.**
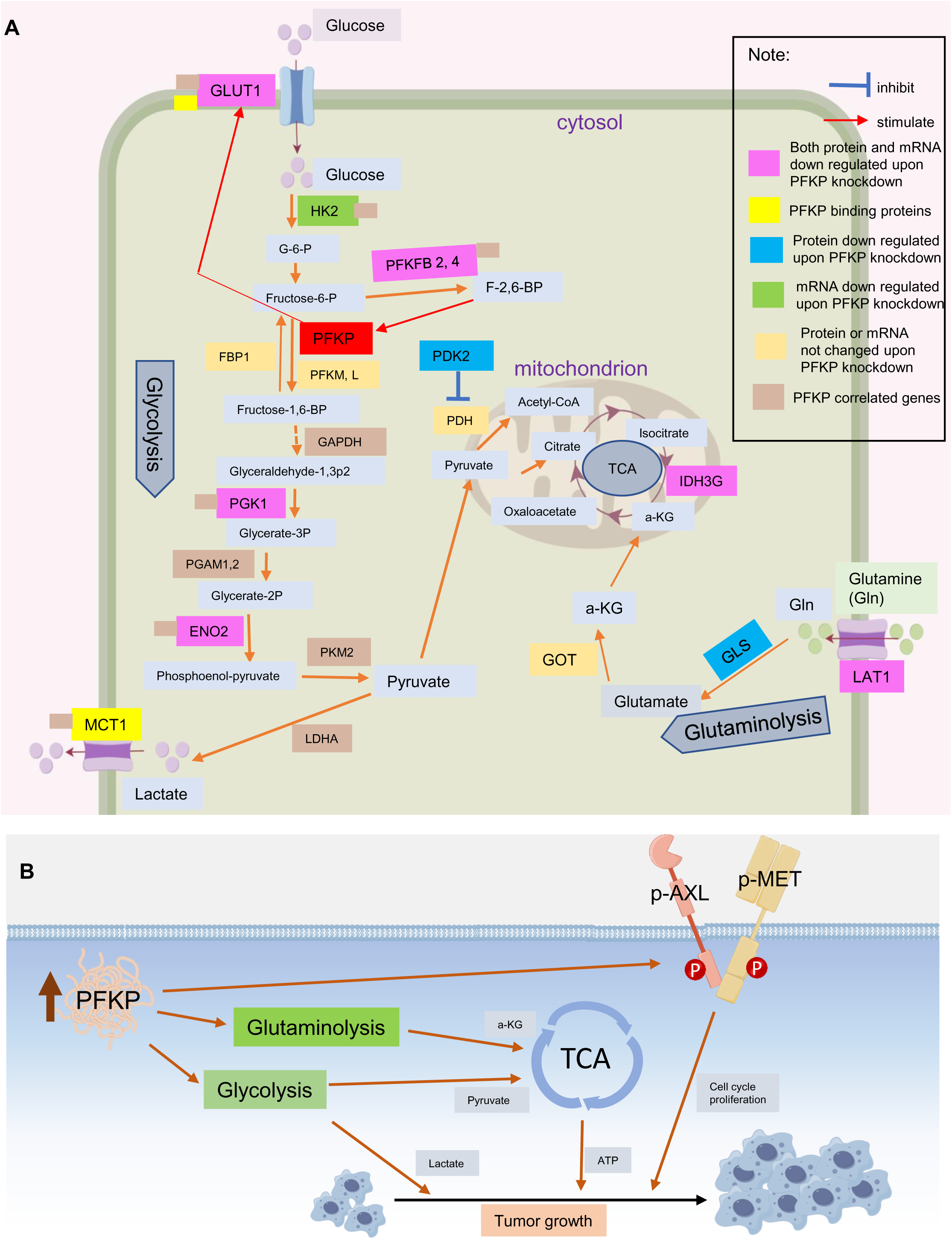
PFKP regulates AXL-MET oncogenic and metabolic pathways in lung cancer. A, DIA-MS, RNA-seq and PFKP Co-IP results indicated that several key proteins (highlighted in purple, blue and green colors) in glycolysis, glutaminolysis and TCA cycle processes were decreased upon PFKP knockdown. GLUT1 and MCT1 may directly bind to PFKP protein (highlighted in yellow color) measured by PFKP Co-IP and MS; **B**, model of the roles of PFKP in lung cancer progression. PFKP is involved in both oncogenic and metabolic pathways in lung cancer progression.

AXL interacting with PFKP was not reported previously, we look at if these PFKP regulated genes/proteins uncovered in this study have potentially protein-protein interaction networks, interestingly we found that they were connected in one network with PFKP, except AXL and MET, using String website (**Figure S7**). In this study, we uncovered that PFKP may directly interact with AXL, suggesting that PFKP plays critical roles in not only metabolic process, but also in non-metabolic pathways for lung cancer progression (**Figure 7B**).

## 4. Discussion

In this study, we demonstrated that the glycolytic enzyme PFKP is highly expressed in multiple types of cancers and significantly contributes to poor patient survival. In addition, we demonstrated a role of PFKP in promoting NSCLC proliferation, migration, invasion and colony formation which together with prior reports[17; 18] confirms that PFKP plays an essential role in lung cancer.

Mechanistically, we found that PFKP has both metabolic and non-metabolic roles in cancer progression. PFKP can directly bind to AXL and promote AXL phosphorylation at Y799, this activation of AXL further leads to phosphorylation of MET through receptor tyrosine kinase hetero-interaction, and the activation of MET and AXL pathways contribute to NSCLC progression. In addition, PFKP can regulate not only glycolysis pathway, but also glutaminolysis and TCA cycle signaling. This is in line with previous reports regarding how metabolic enzymes could affect cell signaling in a non-canonical way[5], and our results provide further evidence for the important role of metabolic enzymes in cell signaling transduction.

PFKP has previously been reported to have roles in several oncogenic signaling pathways including PI3K-AKT[14] and beta-catenin[57]. Further, PFKP undergoes TRIM21-mediated polyubiquitylation and degradation, and this degradation can be abolished either by AKT-mediated stabilization of PFKP[15] or mechanical stress induced TRIM21 sequestering [58], both mechanisms can lead to enhanced glycolysis and tumorigenesis. PFKP can translocate to the nucleus and stimulate CXCR4 expression in T cell malignancy[9]. These results imply that PFKP is a key player not only in cancer cell glucose metabolism, but also in cancer related signaling pathways. AXL is an important therapeutic target in multiple types of cancers[52], and it has important functions in promoting cell proliferation, invasion and migration[59]. AXL activation is also an important mechanism of NSCLC resistance to EGFR inhibitors[51], and targeting both AXL and other RTKs provides a strong response in multiple cancers[60–62]. In addition to canonical ligand-dependent activation, AXL can undergo heterodimerization with other RTKs for its activation, and thus promote activation of the interactor[23–26]. While Y702 phosphorylation is primarily ligand-dependent, the Y779 phosphorylation site is tightly related to AXL heterodimerization with other RTKs. In this study, we found that PFKP binds directly with AXL, and AXL Y779 phosphorylation is decreased upon PFKP knockdown, suggesting that PFKP’s binding promotes AXL phosphorylation at Y779, however, it is still not clear whether PFKP phosphorylates AXL directly or promotes the phosphorylation by other mechanisms. Further *in vitro* kinase assays are needed for further exploration of these events.

AXL undergoes hyper glycosylation modification[63], and in line with previously reports, we found two main protein forms of AXL in NSCLC, one being 120kD and the other at 140kD. It has been reported that the Y779 phosphorylation mainly exist in the 140kD form, and 140kD AXL is the main player in RTK heterodimerization[27]. In our study, we found that the 140kD form AXL also dominates the interaction with PFKP, which is consistent with our hypothesized PFKP-AXL-MET axis mechanism. In addition, as a glycolytic enzyme, PFKP expression is increased in a high glucose environment[64], as does AXL glycosylation levels[63]. Thus, a high glucose concentration, on one hand can promote the expression of PFKP, but also can promote AXL glycosylation, both mechanisms can induce AXL Y779 phosphorylation and subsequent MET activation. This suggests a novel and powerful mechanism for how high glucose levels may contribute to cancer and why cancer cells are more dependent on glucose than normal cells[65]. Taken together, PFKP is an oncogene with significant non-metabolic oncogenic functions, which acts through binding and promoting AXL Y779 phosphorylation, and activating MET by subsequent Y1234/5 phosphorylation, thereby regulating cell proliferation, invasion, migration, and colony formation.

Interestingly, not only several glycolytic proteins such as PGK1, ENO1 and PFKFB4 were affected by PFKP, but also glutaminolytic and TCA cycle enzymes such as GLS and IDH3 may also be regulated by PFKP. Finally, three major transporters, GLUT1, LAT1 and MCT1 may be also regulated by PFKP through either directly binding and/or transcriptional regulation.

## 5. Conclusions

In summary, PFKP is highly expressed in lung cancer and significantly contributes to poor patient survival. PFKP could promote cell proliferation, migration, invasion and colony formation in NSCLC. PFKP plays critical roles in metabolic and non-metabolic pathways for lung cancer progression. PFKP could be a potential novel biomarker for NSCLC. The PFKP-AXL interaction could also be a potential therapeutic target in patients with NSCLC.

## Authorship contributions

YS, HZ, and GC designed experiments. YS, and GC wrote original draft, data curation and data analysis; HF, YL, YL, ZZ, YD, SC, XZ, HR, and WS data correlation and analysis; QM, and GC conceptualization, supervision and funding acquisition.

## Acknowledgments

This work was supported in part by Shenzhen Municipal Science and Technology Innovation Commission Foundation (JCYJ20220530114415036, JCYJ20210324104800001 to G.C.); the National Natural Science Foundation of China (NSFC) (32070625 to G.C., 32100459 to X.Z.). The Center for Computational Science and Engineering of Southern University of Science and Technology.

## Data availability

Data generated in this study are included in the article and raw data will be available upon request.

## Appendix A. Supporting information

Supplementary data associated with this article can be found in the online version

