## Supplemental files for "PFKP regulates AXL-MET oncogenic and metabolic pathways in lung cancer"

**Sun et al. PFKP ms,      Supplementary Tables**

**Table S1 cell lines**

| Cell Line | Subtype | KRAS | TP53 | MET | AXL |
|---|---|---|---|---|---|
| A549 | AD | G12S | WT | WT | WT |
| H1299 | AD | WT | trancated | WT | WT |
| H838 | AD | WT, amp? | E62stop | I638L | WT |

**Table S2 siRNAs**

| Gene name | ID | sequence | company |
|---|---|---|---|
| AXL | AXL-homo-2040 | CCUGUGGUCAUCUUACCUUTT | Genepharma |
| AXL | AXL-homo-1544 | GUGGUAUGUACUGCUAGGATT | Genepharma |
| AXL | AXL-homo-2672 | GGGUGGAGGUUAUCCUGAATT | Genepharma |
| PFKP | PFKP-Homo-1252 | CCGGAUGAUCAGAUCCCAATT | Genepharma |
| PFKP | PFKP-Homo-1367 | GCCACAGGAUGCUCGCCAUTT | Genepharma |
| PFKP | PFKP-Homo-2195 | CCAUUUGUGUGCUGGGAAUTT | Genepharma |

**Table S3 primers**

| gene name | primer ID | primer sequence(5'-3') | product size |
|---|---|---|---|
| PFKP | PFKP R | AAACTCTCGGAGAACCGTGC | 201 |
| PFKP | PFKP F | GAAGGGGTCCCTCCTCTCTG | 201 |
| MET | MET R | GCGATGTTGACATGCCACTG | 216 |
| MET | MET F | TAGCCAACCGAGAGACAAGC | 216 |
| ACTIN | ACTIN R | CTCCTTAATGTCACGCACGAT | 270 |
| ACTIN | ACTIN F | CATGTACGTTGCTATCCAGGC | 270 |

**Table S4 antibodies**

| | |
|---|---|
| p-MET(D26) | Cell Signaling Technology#3077 |
| T-MET(D1C2) | Cell Signaling Technology #8198 |
| PFKP(D2E5) | Cell Signaling Technology #12746 |
| AXL (C89E7) | Cell Signaling Technology#8661 |
| P-702 AXL(D12B2) | Cell Signaling Technology#5724 |
| P-779 AXL | R&D Systems AF2228 |
| β-Actin (8H10D10) | Cell Signaling Technology#12262 |
| FLAG DYKDDDDK Tag (D6W5B) | Cell Signaling Technology#14793 |

| Table S5 56 glycolytic genes mRNA expression | | | |
|---|---|---|---|
| Glycolytic genes | Normal relative expression(log2 PFKP) | Tumor relative expression(log2 PFKP) | Tumor/ Normal ratio (fold change) |
| SLC2A1 | 3.7 | 6.1 | 5.28 |
| PFKP | 4.6 | 6 | 2.64 |
| ENO1 | 9 | 10.3 | 2.46 |
| PC | 3 | 4.3 | 2.46 |
| LDHA | 8.2 | 9.4 | 2.30 |
| GAPDH | 10.2 | 11.3 | 2.14 |
| SLC16A3 | 5.7 | 6.7 | 2.00 |
| TPI1 | 8.4 | 9.3 | 1.87 |
| PKM | 8.5 | 9.4 | 1.87 |
| ALDOA | 9.6 | 10.5 | 1.87 |
| PGK1 | 8 | 8.8 | 1.74 |
| GPI | 6.8 | 7.6 | 1.74 |
| PDK1 | 2.3 | 3 | 1.62 |
| LDHB | 8.1 | 8.7 | 1.52 |
| ALDOC | 3.9 | 4.4 | 1.41 |
| BPGM | 3.7 | 4.2 | 1.41 |
| PGAM1 | 7.1 | 7.4 | 1.23 |
| TIGAR | 2.6 | 2.9 | 1.23 |
| SLC2A11 | 3.2 | 3.5 | 1.23 |
| LDHC | 0.1 | 0.3 | 1.15 |
| PDHX | 3.8 | 4 | 1.15 |
| PDK2 | 4.4 | 4.6 | 1.15 |
| SLC2A14 | 0.2 | 0.3 | 1.07 |
| HK2 | 3.8 | 3.9 | 1.07 |
| PDHB | 5.5 | 5.6 | 1.07 |
| PFKFB1 | 0.2 | 0.2 | 1.00 |
| PDK3 | 2.6 | 2.6 | 1.00 |
| PGAM4 | 0.2 | 0.2 | 1.00 |
| HK1 | 5.8 | 5.7 | 0.93 |
| PCK2 | 4.6 | 4.5 | 0.93 |
| SLC2A10 | 3.1 | 3 | 0.93 |
| ENO4 | 0.4 | 0.3 | 0.93 |
| SLC16A1 | 2.4 | 2.2 | 0.87 |
| PFKL | 6.5 | 6.3 | 0.87 |
| PFKFB4 | 2.6 | 2.4 | 0.87 |
| SLC16A8 | 0.9 | 0.7 | 0.87 |
| ALDOB | 0.4 | 0.2 | 0.87 |
| FBP2 | 0.3 | 0.1 | 0.87 |
| PCK1 | 0.2 | 0 | 0.87 |
| ENO2 | 5.3 | 5 | 0.81 |

| | | | |
|---|---|---|---|
| SLC2A8 | 4.1 | 3.8 | 0.81 |
| SLC16A7 | 2 | 1.7 | 0.81 |
| PGAM2 | 0.8 | 0.5 | 0.81 |
| PDHA1 | 6.6 | 6.2 | 0.76 |
| SLC2A12 | 1.3 | 0.9 | 0.76 |
| GCK | 0.6 | 0.2 | 0.76 |
| PFKM | 4.9 | 4.5 | 0.76 |
| FBP1 | 6.8 | 6.3 | 0.71 |
| SLC2A6 | 3.2 | 2.4 | 0.57 |
| PFKFB3 | 6 | 5.1 | 0.54 |
| PFKFB2 | 4.2 | 3.2 | 0.50 |
| SLC2A9 | 3 | 2 | 0.50 |
| SLC2A4 | 1.4 | 0.3 | 0.47 |
| ENO3 | 4.2 | 2.8 | 0.38 |
| SLC2A3 | 6.5 | 4.1 | 0.19 |
| HK3 | 5.4 | 3 | 0.19 |

**Table S6 Potential interactors of PFKP measured by Co-IP-MS**

| gene name | Protein name | Accession | -10lgP |
|---|---|---|---|
| PFKP | PFKAP | sp\|Q01813\|PFKAP_HUMAN | 297.64 |
| AXL | UFO | sp\|P30530\|UFO_HUMAN | 59.57 |
| SLC2A1 | GTR1 | sp\|P11166\|GTR1_HUMAN | 26.98 |
| SLC16A1 | MOT1 | sp\|P53985\|MOT1_HUMAN | 51.17 |
| ABCE1 | ABCE1 | sp\|P61221\|ABCE1_HUMAN | 81.71 |
| ACIN1 | ACINU | sp\|Q9UKV3\|ACINU_HUMAN | 34.08 |
| AKAP12 | AKA12 | sp\|Q02952\|AKA12_HUMAN | 64.07 |
| AKAP13 | AKP13 | sp\|Q12802\|AKP13_HUMAN | 20.3 |
| AP2M1 | AP2M1 | sp\|Q96CW1\|AP2M1_HUMAN | 20.33 |
| ARF1 | ARF1 | sp\|P84077\|ARF1_HUMAN | 113.02 |
| ARF3 | ARF3 | sp\|P61204\|ARF3_HUMAN | 113.02 |
| ARF4 | ARF4 | sp\|P18085\|ARF4_HUMAN | 96.13 |
| ATAD3A | ATD3A | sp\|Q9NVI7\|ATD3A_HUMAN | 55.66 |
| atad3b | ATD3B | sp\|Q5T9A4\|ATD3B_HUMAN | 55.66 |
| ATP2A2 | AT2A2 | sp\|P16615\|AT2A2_HUMAN | 126.58 |
| ATP5F1C | ATPG | sp\|P36542\|ATPG_HUMAN | 96.18 |
| bag4 | BAG4 | sp\|O95429\|BAG4_HUMAN | 79.28 |
| CALML5 | CALL5 | sp\|Q9NZT1\|CALL5_HUMAN | 29.08 |
| CALR3 | CALR3 | sp\|Q96L12\|CALR3_HUMAN | 29.79 |
| CAND1 | CAND1 | sp\|Q86VP6\|CAND1_HUMAN | 127.86 |
| CAST | ICAL | sp\|P20810\|ICAL_HUMAN | 37.38 |
| CCDC150 | CC150 | sp\|Q8NCX0\|CC150_HUMAN | 22.91 |
| CD59 | CD59 | sp\|P13987\|CD59_HUMAN | 66.14 |
| CDC42 | CDC42 | sp\|P60953\|CDC42_HUMAN | 32.33 |

| | | | |
|---|---|---|---|
| CDK1 | CDK1 | sp|P06493|CDK1_HUMAN | 99.04 |
| CLU | CLUS | sp|P10909|CLUS_HUMAN | 21.48 |
| CNN3 | CNN3 | sp|Q15417|CNN3_HUMAN | 37.89 |
| COX6C | COX6C | sp|P09669|COX6C_HUMAN | 29.64 |
| CSTA | CYTA | sp|P01040|CYTA_HUMAN | 26.6 |
| DDX39A | DX39A | sp|O00148|DX39A_HUMAN | 102.12 |
| DEK | DEK | sp|P35659|DEK_HUMAN | 103.94 |
| DIAPH1 | DIAP1 | sp|O60610|DIAP1_HUMAN | 26.48 |
| DNAJC10 | DJC10 | sp|Q8IXB1|DJC10_HUMAN | 67.47 |
| DNM2 | DYN2 | sp|P50570|DYN2_HUMAN | 53.61 |
| DNM3 | DYN3 | sp|Q9UQ16|DYN3_HUMAN | 53.61 |
| EEF1G | EF1G | sp|P26641|EF1G_HUMAN | 168.64 |
| EHD4 | EHD4 | sp|Q9H223|EHD4_HUMAN | 91.89 |
| EIF3L | EIF3L | sp|Q9Y262|EIF3L_HUMAN | 88.78 |
| EMD | EMD | sp|P50402|EMD_HUMAN | 97.15 |
| FAM168A | F168A | sp|Q92567|F168A_HUMAN | 49.57 |
| FAM83G | FA83G | sp|A6ND36|FA83G_HUMAN | 65.99 |
| FARSA | SYFA | sp|Q9Y285|SYFA_HUMAN | 83.08 |
| FBL | FBRL | sp|P22087|FBRL_HUMAN | 122.05 |
| FHL1 | FHL1 | sp|Q13642|FHL1_HUMAN | 43.58 |
| FHL3 | FHL3 | sp|Q13643|FHL3_HUMAN | 36.71 |
| FLII | FLII | sp|Q13045|FLII_HUMAN | 47.56 |
| FN1 | FINC | sp|P02751|FINC_HUMAN | 45.11 |
| G6PD | G6PD | sp|P11413|G6PD_HUMAN | 25.34 |
| GNL3 | GNL3 | sp|Q9BVP2|GNL3_HUMAN | 64.43 |
| GOT1 | AATC | sp|P17174|AATC_HUMAN | 43.45 |
| H1-4 | H14 | sp|P10412|H14_HUMAN | 150.6 |
| H2AC21 | H2A2B | sp|Q8IUE6|H2A2B_HUMAN | 51.45 |
| H2AC4 | H2A1B | sp|P04908|H2A1B_HUMAN | 65.99 |
| H2AC6 | H2A1C | sp|Q93077|H2A1C_HUMAN | 65.99 |
| H2AW | H2A3 | sp|Q7L7L0|H2A3_HUMAN | 65.99 |
| H2AZ1 | H2AZ | sp|P0C0S5|H2AZ_HUMAN | 69.06 |
| H2AZ2 | H2AV | sp|Q71UI9|H2AV_HUMAN | 69.06 |
| HELLS | HELLS | sp|Q9NRZ9|HELLS_HUMAN | 28.11 |
| HERC5 | HERC5 | sp|Q9UII4|HERC5_HUMAN | 20.16 |
| HIGD1A | HIG1A | sp|Q9Y241|HIG1A_HUMAN | 26.79 |
| HNRNPC | HNRPC | sp|P07910|HNRPC_HUMAN | 94.09 |
| HNRNPH2 | HNRH2 | sp|P55795|HNRH2_HUMAN | 179.46 |
| hoxc8 | HXC8 | sp|P31273|HXC8_HUMAN | 77.95 |
| HRNR | HORN | sp|Q86YZ3|HORN_HUMAN | 106.62 |
| HTRA1 | HTRA1 | sp|Q92743|HTRA1_HUMAN | 24.32 |
| iars2 | SYIM | sp|Q9NSE4|SYIM_HUMAN | 40.89 |
| IGKV2D-26 | KVD26 | sp|A0A0A0MRZ7|KVD26_HUMAN | 125.47 |
| IGKV2D-29 | KVD29 | sp|A0A075B6S2|KVD29_HUMAN | 125.47 |
| KLF4 | KLF4 | sp|O43474|KLF4_HUMAN | 72.18 |

| | | | |
|---|---|---|---|
| KRR1 | KRR1 | sp\|Q13601\|KRR1_HUMAN | 42.69 |
| KRT77 | K2C1B | sp\|Q7Z794\|K2C1B_HUMAN | 127.92 |
| LAS1L | LAS1L | sp\|Q9Y4W2\|LAS1L_HUMAN | 31.25 |
| MACROH2A1 | H2AY | sp\|O75367\|H2AY_HUMAN | 36.19 |
| MAGED2 | MAGD2 | sp\|Q9UNF1\|MAGD2_HUMAN | 131.53 |
| MAP2K3 | MP2K3 | sp\|P46734\|MP2K3_HUMAN | 68.16 |
| MMS19 | MMS19 | sp\|Q96T76\|MMS19_HUMAN | 28.49 |
| MUC19 | MUC19 | sp\|Q7Z5P9\|MUC19_HUMAN | 43.11 |
| NDUFA10 | NDUAA | sp\|O95299\|NDUAA_HUMAN | 46.56 |
| NDUFS2 | NDUS2 | sp\|O75306\|NDUS2_HUMAN | 28.67 |
| NDUFS7 | NDUS7 | sp\|O75251\|NDUS7_HUMAN | 44.21 |
| NOC3L | NOC3L | sp\|Q8WTT2\|NOC3L_HUMAN | 29.26 |
| NOP56 | NOP56 | sp\|O00567\|NOP56_HUMAN | 110.86 |
| NOP58 | NOP58 | sp\|Q9Y2X3\|NOP58_HUMAN | 112.47 |
| NQO1 | NQO1 | sp\|P15559\|NQO1_HUMAN | 27.34 |
| NSA2 | NSA2 | sp\|O95478\|NSA2_HUMAN | 52.7 |
| NSD1 | NSD1 | sp\|Q96L73\|NSD1_HUMAN | 23.84 |
| NUP85 | NUP85 | sp\|Q9BW27\|NUP85_HUMAN | 29.4 |
| NUP93 | NUP93 | sp\|Q8N1F7\|NUP93_HUMAN | 99.99 |
| PABPC4 | PABP4 | sp\|Q13310\|PABP4_HUMAN | 119.42 |
| PABPN1 | PABP2 | sp\|Q86U42\|PABP2_HUMAN | 47.19 |
| PFKL | PFKAL | sp\|P17858\|PFKAL_HUMAN | 163.34 |
| PFKM | PFKAM | sp\|P08237\|PFKAM_HUMAN | 151.83 |
| PGAM5 | PGAM5 | sp\|Q96HS1\|PGAM5_HUMAN | 31.22 |
| pip | PIP | sp\|P12273\|PIP_HUMAN | 33.06 |
| PLBD2 | PLBL2 | sp\|Q8NHP8\|PLBL2_HUMAN | 80.9 |
| PRDX1 | PRDX1 | sp\|Q06830\|PRDX1_HUMAN | 83.93 |
| prss1 | TRY1 | sp\|P07477\|TRY1_HUMAN | 48.92 |
| PSAT1 | SERC | sp\|Q9Y617\|SERC_HUMAN | 50.39 |
| PSMB5 | PSB5 | sp\|P28074\|PSB5_HUMAN | 36.33 |
| PSMD3 | PSMD3 | sp\|O43242\|PSMD3_HUMAN | 89.06 |
| PSMG1 | PSMG1 | sp\|O95456\|PSMG1_HUMAN | 29.55 |
| PXN | PAXI | sp\|P49023\|PAXI_HUMAN | 75.59 |
| QARS1 | SYQ | sp\|P47897\|SYQ_HUMAN | 63.2 |
| **RAB10** | RAB10 | sp\|P61026\|RAB10_HUMAN | 79.18 |
| **RAB13** | RAB13 | sp\|P51153\|RAB13_HUMAN | 41.37 |
| **RAB14** | RAB14 | sp\|P61106\|RAB14_HUMAN | 56.4 |
| **RAB32** | RAB32 | sp\|Q13637\|RAB32_HUMAN | 86.96 |
| **RAB34** | RAB34 | sp\|Q9BZG1\|RAB34_HUMAN | 64.48 |
| **RAB3B** | RAB3B | sp\|P20337\|RAB3B_HUMAN | 83.66 |
| **RAB7A** | RAB7A | sp\|P51149\|RAB7A_HUMAN | 90.09 |
| RBM5 | RBM5 | sp\|P52756\|RBM5_HUMAN | 51.15 |
| RCN1 | RCN1 | sp\|Q15293\|RCN1_HUMAN | 25.29 |
| RPL36A | RL36A | sp\|P83881\|RL36A_HUMAN | 103.07 |
| RPN1 | RPN1 | sp\|P04843\|RPN1_HUMAN | 146.3 |

| | | | |
|---|---|---|---|
| RPS10 | RS10 | sp\|P46783\|RS10_HUMAN | 23.07 |
| RPS27L | RS27L | sp\|Q71UM5\|RS27L_HUMAN | 81.74 |
| RSL1D1 | RL1D1 | sp\|O76021\|RL1D1_HUMAN | 27.94 |
| SAMHD1 | SAMH1 | sp\|Q9Y3Z3\|SAMH1_HUMAN | 23.33 |
| SCD | ACOD | sp\|O00767\|ACOD_HUMAN | 79.01 |
| SEC61A1 | S61A1 | sp\|P61619\|S61A1_HUMAN | 50.65 |
| SEPTIN8 | 8-Sep | sp\|Q92599\|SEPT8_HUMAN | 38.34 |
| SF3A3 | SF3A3 | sp\|Q12874\|SF3A3_HUMAN | 44.4 |
| SFXN1 | SFXN1 | sp\|Q9H9B4\|SFXN1_HUMAN | 71.11 |
| **SLC16A3** | MOT4 | sp\|O15427\|MOT4_HUMAN | 82.64 |
| **SLC25A1** | TXTP | sp\|P53007\|TXTP_HUMAN | 81.16 |
| **SLC25A10** | DIC | sp\|Q9UBX3\|DIC_HUMAN | 50.18 |
| **SLC25A11** | M2OM | sp\|Q02978\|M2OM_HUMAN | 38.47 |
| **SLC25A12** | CMC1 | sp\|O75746\|CMC1_HUMAN | 74.86 |
| **SLC25A13** | CMC2 | sp\|Q9UJS0\|CMC2_HUMAN | 101.97 |
| **SLC25A3** | MPCP | sp\|Q00325\|MPCP_HUMAN | 117.09 |
| **SLC25A5** | ADT2 | sp\|P05141\|ADT2_HUMAN | 146.62 |
| **SLC2A14** | GTR14 | sp\|Q8TDB8\|GTR14_HUMAN | 38.22 |
| SPAG9 | JIP4 | sp\|O60271\|JIP4_HUMAN | 24.32 |
| SPTLC1 | SPTC1 | sp\|O15269\|SPTC1_HUMAN | 39.91 |
| SQOR | SQOR | sp\|Q9Y6N5\|SQOR_HUMAN | 30.33 |
| SRSF1 | SRSF1 | sp\|Q07955\|SRSF1_HUMAN | 49.27 |
| SRSF3 | SRSF3 | sp\|P84103\|SRSF3_HUMAN | 34.74 |
| STAU1 | STAU1 | sp\|O95793\|STAU1_HUMAN | 43.42 |
| SUGP2 | SUGP2 | sp\|Q8IX01\|SUGP2_HUMAN | 48.89 |
| TDRD12 | TDR12 | sp\|Q587J7\|TDR12_HUMAN | 77.12 |
| TECR | TECR | sp\|Q9NZ01\|TECR_HUMAN | 71.24 |
| TIMMDC1 | TIDC1 | sp\|Q9NPL8\|TIDC1_HUMAN | 32.23 |
| TMEM165 | TM165 | sp\|Q9HC07\|TM165_HUMAN | 25.78 |
| TMEM33 | TMM33 | sp\|P57088\|TMM33_HUMAN | 63.2 |
| TOR1AIP1 | TOIP1 | sp\|Q5JTV8\|TOIP1_HUMAN | 25.88 |
| TRA2B | TRA2B | sp\|P62995\|TRA2B_HUMAN | 60.97 |
| TRIP13 | PCH2 | sp\|Q15645\|PCH2_HUMAN | 73.97 |
| TUBB6 | TBB6 | sp\|Q9BUF5\|TBB6_HUMAN | 153.14 |
| U2AF1 | U2AF1 | sp\|Q01081\|U2AF1_HUMAN | 52.76 |
| U2AF1L5 | U2AF5 | sp\|P0DN76\|U2AF5_HUMAN | 52.76 |
| U2AF2 | U2AF2 | sp\|P26368\|U2AF2_HUMAN | 54.58 |
| XPOT | XPOT | sp\|O43592\|XPOT_HUMAN | 56.34 |
| ZC3H14 | ZC3HE | sp\|Q6PJT7\|ZC3HE_HUMAN | 81.41 |
| ZC3HAV1 | ZCCHV | sp\|Q7Z2W4\|ZCCHV_HUMAN | 83.22 |
| ZKSCAN8 | ZKSC8 | sp\|Q15776\|ZKSC8_HUMAN | 62.99 |
| ZNF420 | ZN420 | sp\|Q8TAQ5\|ZN420_HUMAN | 97.74 |
| ZNF668 | ZN668 | sp\|Q96K58\|ZN668_HUMAN | 57.91 |
| ZNF696 | ZN696 | sp\|Q9H7X3\|ZN696_HUMAN | 61.61 |

| | Table S7 PFKP correlated genes in LUAD | | | |
|---|---|---|---|---|
| Dataset | Seo data | Dhanasekaran data | Collisson data | |
| Number of LUAD | 87 | 67 | 312 | 466 |
| gene_name | PFKP r value | PFKP r value | PFKP r value | Mean PFKP r value |
| PFKP | 1.00 | 1.00 | 1.00 | 1.00 |
| SLC2A1 | 0.49 | 0.44 | 0.56 | 0.50 |
| SLC16A3 | 0.61 | 0.48 | 0.34 | 0.48 |
| LDHA | 0.46 | 0.37 | 0.51 | 0.45 |
| ERO1L | 0.50 | 0.37 | 0.45 | 0.44 |
| GAPDH | 0.37 | 0.36 | 0.58 | 0.44 |
| CEP55 | 0.49 | 0.30 | 0.51 | 0.43 |
| MCM10 | 0.39 | 0.40 | 0.49 | 0.43 |
| YME1L1 | 0.43 | 0.47 | 0.39 | 0.43 |
| PGAM1 | 0.45 | 0.36 | 0.46 | 0.42 |
| MKI67 | 0.50 | 0.27 | 0.49 | 0.42 |
| EGLN3 | 0.52 | 0.37 | 0.36 | 0.42 |
| IQGAP3 | 0.39 | 0.39 | 0.47 | 0.41 |
| LOXL2 | 0.58 | 0.32 | 0.34 | 0.41 |
| ANGPTL4 | 0.30 | 0.54 | 0.38 | 0.41 |
| CDCA5 | 0.37 | 0.35 | 0.49 | 0.40 |
| RRM2 | 0.44 | 0.36 | 0.39 | 0.40 |
| AHNAK2 | 0.49 | 0.40 | 0.29 | 0.40 |
| OPTN | 0.51 | 0.26 | 0.41 | 0.39 |
| GTSE1 | 0.40 | 0.42 | 0.36 | 0.39 |
| AK4 | 0.39 | 0.37 | 0.42 | 0.39 |
| NAMPT | 0.50 | 0.24 | 0.43 | 0.39 |
| ARNTL2 | 0.53 | 0.29 | 0.35 | 0.39 |
| NEIL3 | 0.43 | 0.38 | 0.37 | 0.39 |
| GTPBP4 | 0.45 | 0.30 | 0.41 | 0.39 |
| CDCA8 | 0.34 | 0.37 | 0.44 | 0.38 |
| TK1 | 0.36 | 0.42 | 0.37 | 0.38 |
| SPAG5 | 0.41 | 0.34 | 0.39 | 0.38 |
| PGK1 | 0.53 | 0.52 | 0.09 | 0.38 |
| FOXM1 | 0.34 | 0.31 | 0.49 | 0.38 |
| TPI1 | 0.34 | 0.31 | 0.49 | 0.38 |
| DLGAP5 | 0.40 | 0.31 | 0.43 | 0.38 |

| | | | | |
|---|---|---|---|---|
| CDK1 | 0.33 | 0.37 | 0.43 | 0.38 |
| FOSL1 | 0.40 | 0.29 | 0.44 | 0.38 |
| TSKU | 0.43 | 0.38 | 0.32 | 0.38 |
| MELK | 0.45 | 0.28 | 0.39 | 0.38 |
| PKM | 0.60 | 0.41 | 0.12 | 0.37 |
| KIF11 | 0.42 | 0.26 | 0.44 | 0.37 |
| HJURP | 0.40 | 0.34 | 0.38 | 0.37 |
| BIRC5 | 0.36 | 0.42 | 0.34 | 0.37 |
| PBK | 0.33 | 0.41 | 0.38 | 0.37 |
| ARHGAP11A | 0.41 | 0.34 | 0.35 | 0.37 |
| INCENP | 0.34 | 0.38 | 0.38 | 0.37 |
| CHEK1 | 0.38 | 0.25 | 0.47 | 0.37 |
| LRP8 | 0.47 | 0.36 | 0.28 | 0.37 |
| PRR11 | 0.36 | 0.35 | 0.39 | 0.37 |
| CCNB1 | 0.38 | 0.33 | 0.40 | 0.37 |
| TXNRD1 | 0.50 | 0.36 | 0.24 | 0.37 |
| ESPL1 | 0.27 | 0.44 | 0.38 | 0.36 |
| ORC1 | 0.36 | 0.33 | 0.40 | 0.36 |
| RBM17 | 0.33 | 0.30 | 0.46 | 0.36 |
| KIF2C | 0.36 | 0.29 | 0.44 | 0.36 |
| GSG2 | 0.44 | 0.30 | 0.36 | 0.36 |
| C15orf48 | 0.51 | 0.30 | 0.28 | 0.36 |
| PLK1 | 0.39 | 0.27 | 0.42 | 0.36 |
| SHCBP1 | 0.36 | 0.34 | 0.39 | 0.36 |
| NAMPTL | 0.47 | 0.18 | 0.43 | 0.36 |
| STC1 | 0.45 | 0.24 | 0.39 | 0.36 |
| MTL5 | 0.28 | 0.40 | 0.40 | 0.36 |
| PLAU | 0.51 | 0.22 | 0.35 | 0.36 |
| CDC123 | 0.45 | 0.32 | 0.31 | 0.36 |
| CENPA | 0.39 | 0.34 | 0.36 | 0.36 |
| KIF4A | 0.38 | 0.31 | 0.39 | 0.36 |
| DIAPH3 | 0.40 | 0.31 | 0.36 | 0.36 |
| RHOV | 0.36 | 0.37 | 0.34 | 0.36 |
| FAM64A | 0.38 | 0.35 | 0.34 | 0.36 |
| SLC7A1 | 0.52 | 0.28 | 0.28 | 0.36 |
| MAD2L1 | 0.36 | 0.36 | 0.34 | 0.36 |
| XPNPEP1 | 0.49 | 0.24 | 0.34 | 0.36 |
| ANLN | 0.39 | 0.27 | 0.41 | 0.36 |
| C6orf223 | 0.43 | 0.39 | 0.24 | 0.36 |
| KIF18B | 0.37 | 0.36 | 0.34 | 0.36 |
| FEN1 | 0.26 | 0.34 | 0.47 | 0.36 |
| CDCA3 | 0.24 | 0.37 | 0.46 | 0.35 |
| SLC39A1 | 0.41 | 0.40 | 0.25 | 0.35 |
| TIMM23B | 0.28 | 0.48 | 0.30 | 0.35 |
| RHOF | 0.29 | 0.41 | 0.36 | 0.35 |

| | | | | |
|---|---|---|---|---|
| GALNT2 | 0.33 | 0.42 | 0.31 | 0.35 |
| CCNE1 | 0.41 | 0.33 | 0.32 | 0.35 |
| LRRC42 | 0.39 | 0.35 | 0.32 | 0.35 |
| ZWINT | 0.31 | 0.31 | 0.43 | 0.35 |
| SLCO4A1 | 0.29 | 0.43 | 0.33 | 0.35 |
| RACGAP1 | 0.34 | 0.37 | 0.35 | 0.35 |
| NCAPH | 0.41 | 0.28 | 0.37 | 0.35 |
| S100A16 | 0.40 | 0.36 | 0.29 | 0.35 |
| CDCA2 | 0.34 | 0.34 | 0.37 | 0.35 |
| RAD51 | 0.36 | 0.32 | 0.36 | 0.35 |
| SLC2A5 | 0.47 | 0.24 | 0.34 | 0.35 |
| MFSD12 | 0.43 | 0.35 | 0.27 | 0.35 |
| CDC20 | 0.26 | 0.33 | 0.45 | 0.35 |
| AVL9 | 0.41 | 0.46 | 0.18 | 0.35 |
| PLIN3 | 0.44 | 0.40 | 0.20 | 0.35 |
| KIF14 | 0.35 | 0.26 | 0.43 | 0.35 |
| CKAP2L | 0.41 | 0.25 | 0.38 | 0.35 |
| CCNA2 | 0.34 | 0.29 | 0.40 | 0.35 |
| FAM83A | 0.25 | 0.41 | 0.37 | 0.35 |
| PTER | 0.33 | 0.35 | 0.35 | 0.34 |
| PGM2 | 0.42 | 0.47 | 0.14 | 0.34 |
| P4HA1 | 0.34 | 0.29 | 0.40 | 0.34 |
| CDKN3 | 0.32 | 0.30 | 0.41 | 0.34 |
| DEPDC1B | 0.29 | 0.37 | 0.36 | 0.34 |
| E2F8 | 0.35 | 0.31 | 0.37 | 0.34 |
| PITX1 | 0.31 | 0.33 | 0.38 | 0.34 |
| CENPI | 0.36 | 0.27 | 0.38 | 0.34 |
| ITGB1 | 0.36 | 0.27 | 0.39 | 0.34 |
| FAM40B | 0.45 | 0.20 | 0.37 | 0.34 |
| TRIP13 | 0.36 | 0.29 | 0.37 | 0.34 |
| ECT2 | 0.40 | 0.29 | 0.33 | 0.34 |
| RPE | 0.40 | 0.42 | 0.20 | 0.34 |
| VANGL1 | 0.39 | 0.34 | 0.29 | 0.34 |
| LRFN4 | 0.36 | 0.24 | 0.42 | 0.34 |
| STIL | 0.35 | 0.27 | 0.39 | 0.34 |
| NCAPG2 | 0.48 | 0.20 | 0.33 | 0.34 |
| BUB1B | 0.40 | 0.25 | 0.36 | 0.34 |
| CCNB2 | 0.36 | 0.28 | 0.37 | 0.34 |
| SPC24 | 0.28 | 0.41 | 0.31 | 0.34 |
| ACBD5 | 0.34 | 0.37 | 0.29 | 0.33 |
| MASTL | 0.35 | 0.23 | 0.42 | 0.33 |
| EXO1 | 0.31 | 0.25 | 0.45 | 0.33 |
| GCLM | 0.46 | 0.26 | 0.28 | 0.33 |
| LMNB2 | 0.41 | 0.29 | 0.30 | 0.33 |
| SKA1 | 0.35 | 0.26 | 0.40 | 0.33 |

| | | | | |
|---|---|---|---|---|
| BUB3 | 0.41 | 0.19 | 0.40 | 0.33 |
| PRC1 | 0.33 | 0.28 | 0.39 | 0.33 |
| CCRN4L | 0.23 | 0.49 | 0.28 | 0.33 |
| BCAR3 | 0.24 | 0.44 | 0.32 | 0.33 |
| KPNA2 | 0.40 | 0.26 | 0.33 | 0.33 |
| STC2 | 0.28 | 0.40 | 0.32 | 0.33 |
| DEPDC1 | 0.34 | 0.23 | 0.42 | 0.33 |
| PDE10A | 0.44 | 0.32 | 0.24 | 0.33 |
| PRTFDC1 | 0.33 | 0.31 | 0.35 | 0.33 |
| CDC45 | 0.31 | 0.35 | 0.33 | 0.33 |
| RPS6KA4 | 0.35 | 0.33 | 0.30 | 0.33 |
| KIAA0101 | 0.33 | 0.39 | 0.26 | 0.33 |
| IL15RA | 0.38 | 0.16 | 0.44 | 0.33 |
| C17orf53 | 0.31 | 0.32 | 0.35 | 0.33 |
| HSPA14 | 0.42 | 0.32 | 0.25 | 0.33 |
| CDC6 | 0.32 | 0.31 | 0.35 | 0.33 |
| CDCP1 | 0.47 | 0.23 | 0.28 | 0.33 |
| GCLC | 0.40 | 0.36 | 0.22 | 0.33 |
| STYK1 | 0.43 | 0.27 | 0.28 | 0.33 |
| TROAP | 0.25 | 0.37 | 0.35 | 0.33 |
| MTPAP | 0.28 | 0.36 | 0.35 | 0.33 |
| PLK4 | 0.35 | 0.26 | 0.36 | 0.33 |
| BDKRB1 | 0.25 | 0.48 | 0.25 | 0.33 |
| DDIT4 | 0.32 | 0.37 | 0.29 | 0.32 |
| PSMA5 | 0.36 | 0.39 | 0.23 | 0.32 |
| IRAK1 | 0.55 | 0.31 | 0.12 | 0.32 |
| GPRIN1 | 0.38 | 0.35 | 0.24 | 0.32 |
| CENPK | 0.36 | 0.30 | 0.31 | 0.32 |
| BUB1 | 0.37 | 0.24 | 0.36 | 0.32 |
| CUL2 | 0.34 | 0.30 | 0.32 | 0.32 |
| SEMA4B | 0.48 | 0.27 | 0.22 | 0.32 |
| C10orf55 | 0.47 | 0.23 | 0.26 | 0.32 |
| SFN | 0.40 | 0.33 | 0.24 | 0.32 |
| ZDHHC18 | 0.39 | 0.33 | 0.24 | 0.32 |
| ASF1B | 0.32 | 0.32 | 0.33 | 0.32 |
| NCAPG | 0.33 | 0.24 | 0.39 | 0.32 |
| MTHFD2 | 0.47 | 0.22 | 0.27 | 0.32 |
| TMEM171 | 0.38 | 0.30 | 0.28 | 0.32 |
| HTATIP2 | 0.35 | 0.33 | 0.28 | 0.32 |
| SLC7A11 | 0.45 | 0.29 | 0.22 | 0.32 |
| RTCA | 0.35 | 0.26 | 0.35 | 0.32 |
| PLAUR | 0.55 | 0.15 | 0.26 | 0.32 |
| TPX2 | 0.35 | 0.21 | 0.39 | 0.32 |
| PNP | 0.37 | 0.29 | 0.29 | 0.32 |
| FADD | 0.28 | 0.36 | 0.32 | 0.32 |

| | | | | |
|---|---|---|---|---|
| FRMD8 | 0.30 | 0.35 | 0.30 | 0.32 |
| DHX37 | 0.36 | 0.30 | 0.30 | 0.32 |
| UBE2S | 0.29 | 0.33 | 0.33 | 0.32 |
| YKT6 | 0.37 | 0.50 | 0.08 | 0.32 |
| LRRC59 | 0.36 | 0.45 | 0.14 | 0.32 |
| PPIF | 0.32 | 0.21 | 0.42 | 0.32 |
| TAP2 | 0.45 | 0.29 | 0.20 | 0.32 |
| RGS20 | 0.30 | 0.37 | 0.27 | 0.32 |
| TEAD4 | 0.29 | 0.26 | 0.40 | 0.31 |
| PLOD2 | 0.44 | 0.17 | 0.34 | 0.31 |
| RAD54L | 0.29 | 0.25 | 0.40 | 0.31 |
| CLSPN | 0.35 | 0.20 | 0.39 | 0.31 |
| DSG2 | 0.39 | 0.35 | 0.19 | 0.31 |
| MMP14 | 0.43 | 0.26 | 0.25 | 0.31 |
| FAM72D | 0.27 | 0.21 | 0.45 | 0.31 |
| PRSS23 | 0.29 | 0.41 | 0.24 | 0.31 |
| SEC23A | 0.33 | 0.33 | 0.28 | 0.31 |
| SFXN1 | 0.36 | 0.34 | 0.23 | 0.31 |
| FAM83D | 0.27 | 0.30 | 0.36 | 0.31 |
| PARD3 | 0.35 | 0.22 | 0.36 | 0.31 |
| CFL1 | 0.26 | 0.23 | 0.44 | 0.31 |
| MYBL2 | 0.30 | 0.25 | 0.38 | 0.31 |
| SAPCD2 | 0.25 | 0.33 | 0.35 | 0.31 |
| NUF2 | 0.29 | 0.24 | 0.40 | 0.31 |
| CASC5 | 0.41 | 0.15 | 0.36 | 0.31 |
| HMMR | 0.30 | 0.25 | 0.37 | 0.31 |
| SMS | 0.31 | 0.28 | 0.34 | 0.31 |
| APOL2 | 0.35 | 0.39 | 0.18 | 0.31 |
| NEK2 | 0.25 | 0.25 | 0.43 | 0.31 |
| FGFBP1 | 0.37 | 0.34 | 0.21 | 0.31 |
| AURKB | 0.31 | 0.28 | 0.33 | 0.31 |
| NUSAP1 | 0.37 | 0.24 | 0.31 | 0.31 |
| MYEOV | 0.26 | 0.24 | 0.43 | 0.31 |
| NDC80 | 0.29 | 0.30 | 0.32 | 0.31 |
| GPI | 0.40 | 0.35 | 0.17 | 0.31 |
| SPHK1 | 0.26 | 0.28 | 0.38 | 0.31 |
| TACC3 | 0.26 | 0.34 | 0.32 | 0.31 |
| SLC3A2 | 0.20 | 0.34 | 0.37 | 0.30 |
| FKBP4 | 0.32 | 0.27 | 0.33 | 0.30 |
| S100A6 | 0.36 | 0.43 | 0.13 | 0.30 |
| ST3GAL4 | 0.30 | 0.34 | 0.27 | 0.30 |
| PIP4K2A | 0.36 | 0.18 | 0.38 | 0.30 |
| ASPM | 0.25 | 0.22 | 0.44 | 0.30 |
| E2F2 | 0.37 | 0.21 | 0.33 | 0.30 |
| LRP10 | 0.36 | 0.44 | 0.11 | 0.30 |

| | | | | |
|---|---|---|---|---|
| SEPHS1 | 0.36 | 0.27 | 0.28 | 0.30 |
| DCUN1D5 | 0.31 | 0.28 | 0.32 | 0.30 |
| UBE2C | 0.34 | 0.23 | 0.34 | 0.30 |
| SH3GLB1 | 0.35 | 0.27 | 0.28 | 0.30 |
| KIFC1 | 0.33 | 0.28 | 0.29 | 0.30 |
| FANCI | 0.34 | 0.27 | 0.30 | 0.30 |
| BMS1 | 0.31 | 0.31 | 0.28 | 0.30 |
| CENPF | 0.29 | 0.21 | 0.41 | 0.30 |
| KLF16 | 0.33 | 0.31 | 0.26 | 0.30 |
| SLC35E4 | 0.39 | 0.39 | 0.11 | 0.30 |
| SRPX2 | 0.46 | 0.25 | 0.19 | 0.30 |
| B3GNT5 | 0.34 | 0.23 | 0.33 | 0.30 |
| KIF20A | 0.38 | 0.20 | 0.32 | 0.30 |
| RCE1 | 0.12 | 0.45 | 0.33 | 0.30 |
| C5orf65 | 0.47 | 0.19 | 0.23 | 0.30 |
| RIPK2 | 0.43 | 0.20 | 0.27 | 0.30 |
| ARHGAP11B | 0.45 | 0.26 | 0.19 | 0.30 |
| HK2 | 0.30 | 0.30 | 0.30 | 0.30 |
| ESCO2 | 0.33 | 0.29 | 0.28 | 0.30 |
| LARP1 | 0.36 | 0.42 | 0.11 | 0.30 |
| SBNO2 | 0.40 | 0.32 | 0.17 | 0.30 |
| C9orf40 | 0.35 | 0.25 | 0.29 | 0.30 |
| ADM | 0.33 | 0.19 | 0.37 | 0.30 |
| GJB3 | 0.30 | 0.25 | 0.34 | 0.30 |
| AP5B1 | 0.29 | 0.38 | 0.22 | 0.30 |
| CDT1 | 0.31 | 0.23 | 0.36 | 0.30 |
| CDC25C | 0.27 | 0.25 | 0.36 | 0.30 |
| EIF2S1 | 0.37 | 0.38 | 0.14 | 0.30 |
| RELT | 0.40 | 0.18 | 0.32 | 0.30 |
| CENPM | 0.28 | 0.34 | 0.27 | 0.30 |
| H2AFZ | 0.38 | 0.32 | 0.19 | 0.30 |
| MTFP1 | 0.21 | 0.55 | 0.12 | 0.30 |
| STAM | 0.32 | 0.19 | 0.38 | 0.30 |
| FAM72B | 0.23 | 0.24 | 0.42 | 0.30 |
| NCAPD2 | 0.22 | 0.31 | 0.36 | 0.30 |
| BDKRB2 | 0.36 | 0.40 | 0.12 | 0.29 |
| TAP1 | 0.46 | 0.28 | 0.14 | 0.29 |
| GDI2 | 0.49 | 0.06 | 0.33 | 0.29 |
| AC108463.3 | 0.38 | 0.36 | 0.14 | 0.29 |
| DUSP5 | 0.33 | 0.22 | 0.34 | 0.29 |
| MC1R | 0.25 | 0.46 | 0.17 | 0.29 |
| GINS3 | 0.26 | 0.42 | 0.20 | 0.29 |
| UBE2Z | 0.34 | 0.49 | 0.05 | 0.29 |
| TUBA1B | 0.21 | 0.27 | 0.40 | 0.29 |
| KRT80 | 0.24 | 0.36 | 0.28 | 0.29 |

| | | | | |
|---|---|---|---|---|
| MTHFD1L | 0.34 | 0.32 | 0.22 | 0.29 |
| FOSL2 | 0.49 | 0.28 | 0.11 | 0.29 |
| OIP5 | 0.27 | 0.35 | 0.26 | 0.29 |
| H2AFX | 0.20 | 0.30 | 0.37 | 0.29 |
| HCCS | 0.36 | 0.42 | 0.09 | 0.29 |
| TMEM48 | 0.30 | 0.25 | 0.33 | 0.29 |
| GPC6 | 0.33 | 0.24 | 0.30 | 0.29 |
| C1QTNF6 | 0.29 | 0.35 | 0.24 | 0.29 |
| CENPE | 0.30 | 0.22 | 0.35 | 0.29 |
| ENO1 | 0.21 | 0.36 | 0.30 | 0.29 |
| APOL6 | 0.41 | 0.35 | 0.12 | 0.29 |
| HTR1D | 0.33 | 0.26 | 0.28 | 0.29 |
| CENPL | 0.30 | 0.31 | 0.26 | 0.29 |
| AURKA | 0.22 | 0.28 | 0.37 | 0.29 |
| CYCS | 0.22 | 0.46 | 0.19 | 0.29 |
| HMGA1 | 0.32 | 0.27 | 0.28 | 0.29 |
| CCL26 | 0.35 | 0.25 | 0.27 | 0.29 |
| KIF23 | 0.28 | 0.23 | 0.35 | 0.29 |
| CTSL2 | 0.22 | 0.33 | 0.31 | 0.29 |
| DBF4 | 0.39 | 0.10 | 0.38 | 0.29 |
| UCK2 | 0.24 | 0.21 | 0.41 | 0.29 |
| RAB35 | 0.43 | 0.20 | 0.23 | 0.29 |
| ANKRD16 | 0.39 | 0.29 | 0.18 | 0.29 |
| PGM1 | 0.15 | 0.36 | 0.35 | 0.29 |
| BRI3BP | 0.27 | 0.33 | 0.26 | 0.29 |
| TPM3 | 0.39 | 0.06 | 0.41 | 0.29 |
| SULF1 | 0.50 | 0.16 | 0.20 | 0.29 |
| AACS | 0.31 | 0.35 | 0.20 | 0.29 |
| FAM54A | 0.26 | 0.25 | 0.35 | 0.29 |
| TYMP | 0.38 | 0.37 | 0.12 | 0.29 |
| PKP3 | 0.15 | 0.37 | 0.34 | 0.29 |
| RPL39L | 0.33 | 0.38 | 0.16 | 0.29 |
| MAFG | 0.45 | 0.31 | 0.10 | 0.29 |
| WDR67 | 0.25 | 0.46 | 0.14 | 0.29 |
| C20orf24 | 0.34 | 0.41 | 0.11 | 0.29 |
| LMNB1 | 0.36 | 0.24 | 0.26 | 0.29 |
| VDAC1 | 0.29 | 0.35 | 0.22 | 0.29 |
| PSMD2 | 0.33 | 0.34 | 0.18 | 0.28 |
| BRCA1 | 0.34 | 0.20 | 0.31 | 0.28 |
| S100A10 | 0.26 | 0.32 | 0.28 | 0.28 |
| POC1A | 0.28 | 0.25 | 0.33 | 0.28 |
| SUV39H2 | 0.36 | 0.17 | 0.31 | 0.28 |
| COL11A1 | 0.37 | 0.19 | 0.29 | 0.28 |
| HDGF | 0.28 | 0.24 | 0.34 | 0.28 |
| SPRED3 | 0.24 | 0.40 | 0.21 | 0.28 |

| | | | | |
|---|---|---|---|---|
| LAMC2 | 0.36 | 0.15 | 0.34 | 0.28 |
| MCM6 | 0.33 | 0.25 | 0.27 | 0.28 |
| ERLIN1 | 0.26 | 0.26 | 0.33 | 0.28 |
| VEGFC | 0.29 | 0.32 | 0.24 | 0.28 |
| SKA3 | 0.33 | 0.13 | 0.39 | 0.28 |
| KIAA1524 | 0.30 | 0.22 | 0.33 | 0.28 |
| RAD51AP1 | 0.21 | 0.28 | 0.36 | 0.28 |
| STX1A | 0.41 | 0.25 | 0.19 | 0.28 |
| AP1S3 | 0.30 | 0.31 | 0.24 | 0.28 |
| MYO19 | 0.22 | 0.47 | 0.16 | 0.28 |
| KIF20B | 0.37 | 0.09 | 0.39 | 0.28 |
| ADAM12 | 0.45 | 0.07 | 0.33 | 0.28 |
| ARF6 | 0.39 | 0.28 | 0.16 | 0.28 |
| TUBB3 | 0.40 | 0.14 | 0.31 | 0.28 |
| HNRNPF | 0.20 | 0.36 | 0.27 | 0.28 |
| AKAP12 | 0.20 | 0.36 | 0.28 | 0.28 |
| PPP1R14B | 0.16 | 0.36 | 0.33 | 0.28 |
| KYNU | 0.35 | 0.26 | 0.23 | 0.28 |
| LTBR | 0.22 | 0.34 | 0.27 | 0.28 |
| PMAIP1 | 0.34 | 0.24 | 0.26 | 0.28 |
| RCN1 | 0.42 | 0.24 | 0.18 | 0.28 |
| ABI1 | 0.35 | 0.27 | 0.22 | 0.28 |
| AGFG1 | 0.37 | 0.27 | 0.19 | 0.28 |
| GINS4 | 0.34 | 0.26 | 0.24 | 0.28 |
| CD274 | 0.47 | 0.10 | 0.26 | 0.28 |
| CTHRC1 | 0.43 | 0.24 | 0.16 | 0.28 |
| PHLDA2 | 0.20 | 0.29 | 0.35 | 0.28 |
| SLC12A8 | 0.47 | 0.16 | 0.21 | 0.28 |
| JMJD6 | 0.31 | 0.35 | 0.17 | 0.28 |
| TCF19 | 0.38 | 0.27 | 0.19 | 0.28 |
| HAPLN3 | 0.49 | 0.16 | 0.19 | 0.28 |
| ALDOA | 0.36 | 0.35 | 0.11 | 0.28 |
| LYPD3 | 0.25 | 0.34 | 0.24 | 0.28 |
| SPC25 | 0.31 | 0.27 | 0.24 | 0.28 |
| ZNF367 | 0.33 | 0.21 | 0.29 | 0.28 |
| POLQ | 0.35 | 0.21 | 0.27 | 0.28 |
| PSME2 | 0.34 | 0.31 | 0.18 | 0.28 |
| APOL1 | 0.26 | 0.38 | 0.19 | 0.28 |
| RP11-831H9.16 | 0.34 | 0.30 | 0.19 | 0.28 |
| GAL | 0.40 | 0.14 | 0.29 | 0.27 |
| CENPW | 0.27 | 0.23 | 0.33 | 0.27 |
| POP1 | 0.22 | 0.36 | 0.24 | 0.27 |
| ZC3HAV1L | 0.36 | 0.31 | 0.15 | 0.27 |
| TYMS | 0.30 | 0.23 | 0.29 | 0.27 |
| TRIM16 | 0.48 | 0.25 | 0.09 | 0.27 |

| | | | | |
|---|---|---|---|---|
| MFI2 | 0.27 | 0.24 | 0.32 | 0.27 |
| MCM4 | 0.31 | 0.23 | 0.27 | 0.27 |
| PPP1R3G | 0.17 | 0.35 | 0.30 | 0.27 |
| BIRC3 | 0.32 | 0.27 | 0.23 | 0.27 |
| MOCOS | 0.38 | 0.20 | 0.24 | 0.27 |
| RHBDF2 | 0.41 | 0.22 | 0.19 | 0.27 |
| PPP2R2C | 0.24 | 0.43 | 0.15 | 0.27 |
| PTPRH | 0.17 | 0.32 | 0.33 | 0.27 |
| VOPP1 | 0.35 | 0.20 | 0.27 | 0.27 |
| PPTC7 | 0.12 | 0.50 | 0.20 | 0.27 |
| FAM83B | 0.26 | 0.34 | 0.22 | 0.27 |
| WISP1 | 0.48 | 0.17 | 0.16 | 0.27 |
| AC112721.1 | 0.34 | 0.34 | 0.13 | 0.27 |
| ABCA12 | 0.34 | 0.26 | 0.21 | 0.27 |
| GBP5 | 0.41 | 0.13 | 0.27 | 0.27 |
| PDSS1 | 0.19 | 0.24 | 0.39 | 0.27 |
| P4HA2 | 0.35 | 0.31 | 0.16 | 0.27 |
| ATG16L1 | 0.38 | 0.35 | 0.08 | 0.27 |
| IL4I1 | 0.47 | 0.17 | 0.17 | 0.27 |
| RBM15 | 0.26 | 0.37 | 0.18 | 0.27 |
| PLCB3 | 0.26 | 0.20 | 0.34 | 0.27 |
| HN1 | 0.27 | 0.35 | 0.19 | 0.27 |
| GJB2 | 0.28 | 0.25 | 0.28 | 0.27 |
| SLC20A1 | 0.41 | 0.18 | 0.22 | 0.27 |
| TUFT1 | 0.29 | 0.24 | 0.28 | 0.27 |
| GLT25D1 | 0.40 | 0.20 | 0.20 | 0.27 |
| PSRC1 | 0.14 | 0.36 | 0.31 | 0.27 |
| PLXNA1 | 0.32 | 0.33 | 0.15 | 0.27 |
| SVIL | 0.16 | 0.50 | 0.14 | 0.27 |
| KRT16 | 0.24 | 0.34 | 0.22 | 0.27 |
| C14orf80 | 0.18 | 0.34 | 0.28 | 0.27 |
| RP13-672B3.2 | 0.27 | 0.28 | 0.25 | 0.27 |
| PTPRJ | 0.38 | 0.31 | 0.12 | 0.27 |
| PPAPDC1A | 0.32 | 0.16 | 0.33 | 0.27 |
| FAM72A | 0.18 | 0.19 | 0.44 | 0.27 |
| FBXO45 | 0.24 | 0.30 | 0.26 | 0.27 |
| FAM210A | 0.34 | 0.29 | 0.18 | 0.27 |
| GBP1 | 0.41 | 0.11 | 0.28 | 0.27 |
| GOLGA3 | 0.28 | 0.38 | 0.14 | 0.27 |
| PIF1 | 0.33 | 0.28 | 0.20 | 0.27 |
| CD109 | 0.24 | 0.25 | 0.31 | 0.27 |
| PLEK2 | 0.36 | 0.20 | 0.24 | 0.27 |
| DTL | 0.27 | 0.10 | 0.44 | 0.27 |
| AVEN | 0.46 | 0.15 | 0.20 | 0.27 |
| PFKFB4 | 0.31 | 0.20 | 0.30 | 0.27 |

| | | | | |
|---|---|---|---|---|
| SSX2IP | 0.34 | 0.13 | 0.33 | 0.27 |
| BCL10 | 0.30 | 0.25 | 0.25 | 0.27 |
| CKS1B | 0.18 | 0.29 | 0.33 | 0.27 |
| SELRC1 | 0.26 | 0.33 | 0.21 | 0.27 |
| C19orf12 | 0.47 | 0.31 | 0.02 | 0.27 |
| TDP1 | 0.39 | 0.24 | 0.16 | 0.27 |
| ELK1 | 0.27 | 0.49 | 0.03 | 0.27 |
| SPOCD1 | 0.30 | 0.14 | 0.35 | 0.27 |
| NLN | 0.39 | 0.18 | 0.23 | 0.27 |
| WDR76 | 0.42 | 0.10 | 0.28 | 0.27 |
| ADAR | 0.32 | 0.25 | 0.22 | 0.27 |
| SRXN1 | 0.37 | 0.31 | 0.11 | 0.27 |
| UBE2T | 0.27 | 0.14 | 0.38 | 0.27 |
| DNAJC9 | 0.22 | 0.18 | 0.40 | 0.26 |
| GALNT6 | 0.40 | 0.20 | 0.20 | 0.26 |
| RHPN2 | 0.18 | 0.41 | 0.20 | 0.26 |
| AIM1L | 0.13 | 0.40 | 0.26 | 0.26 |
| TTF2 | 0.25 | 0.30 | 0.24 | 0.26 |
| KLHDC7B | 0.35 | 0.30 | 0.15 | 0.26 |
| ITGA5 | 0.27 | 0.19 | 0.33 | 0.26 |
| RHOC | 0.30 | 0.26 | 0.24 | 0.26 |
| EPSTI1 | 0.39 | 0.22 | 0.19 | 0.26 |
| TICAM1 | 0.25 | 0.31 | 0.22 | 0.26 |
| PPAT | 0.38 | 0.19 | 0.23 | 0.26 |
| GNAI3 | 0.28 | 0.25 | 0.27 | 0.26 |
| DDX21 | 0.34 | 0.14 | 0.31 | 0.26 |
| HHIPL2 | 0.34 | 0.17 | 0.28 | 0.26 |
| DNAJC10 | 0.51 | 0.23 | 0.05 | 0.26 |
| AKIP1 | 0.30 | 0.30 | 0.18 | 0.26 |
| ACTR3 | 0.47 | 0.26 | 0.06 | 0.26 |
| ANKRD32 | 0.35 | 0.18 | 0.26 | 0.26 |
| SOD2 | 0.35 | 0.18 | 0.25 | 0.26 |
| PAICS | 0.37 | 0.23 | 0.18 | 0.26 |
| VEGFA | 0.45 | 0.22 | 0.11 | 0.26 |
| SEC14L2 | 0.20 | 0.42 | 0.16 | 0.26 |
| MCF2L2 | 0.26 | 0.22 | 0.31 | 0.26 |
| IPPK | 0.38 | 0.24 | 0.16 | 0.26 |
| RALA | 0.29 | 0.33 | 0.16 | 0.26 |
| SHOX2 | 0.24 | 0.39 | 0.15 | 0.26 |
| RAVER1 | 0.37 | 0.38 | 0.03 | 0.26 |
| ATP8B3 | 0.28 | 0.26 | 0.25 | 0.26 |
| KCMF1 | 0.37 | 0.31 | 0.10 | 0.26 |
| PGAM5 | 0.23 | 0.30 | 0.25 | 0.26 |
| SLC9A7 | 0.27 | 0.29 | 0.22 | 0.26 |
| KLC2 | 0.16 | 0.29 | 0.33 | 0.26 |

| | | | | |
|---|---|---|---|---|
| FAM189B | 0.15 | 0.31 | 0.31 | 0.26 |
| NMT2 | 0.27 | 0.28 | 0.23 | 0.26 |
| PTGES | 0.30 | 0.35 | 0.13 | 0.26 |
| TOP2A | 0.22 | 0.27 | 0.28 | 0.26 |
| GPR97 | 0.23 | 0.21 | 0.34 | 0.26 |
| OAS3 | 0.33 | 0.17 | 0.28 | 0.26 |
| MCMBP | 0.42 | 0.05 | 0.31 | 0.26 |
| GPR115 | 0.22 | 0.28 | 0.27 | 0.26 |
| PNMA1 | 0.32 | 0.24 | 0.21 | 0.26 |
| MB21D1 | 0.45 | 0.06 | 0.26 | 0.26 |
| PGAM4 | 0.26 | 0.13 | 0.38 | 0.26 |
| KIF18A | 0.22 | 0.19 | 0.36 | 0.26 |
| GBE1 | 0.37 | 0.22 | 0.18 | 0.26 |
| WDHD1 | 0.28 | 0.20 | 0.29 | 0.26 |
| PYGL | 0.19 | 0.26 | 0.33 | 0.26 |
| IRAK2 | 0.41 | 0.21 | 0.14 | 0.26 |
| BRIP1 | 0.30 | 0.19 | 0.28 | 0.26 |
| ABCE1 | 0.28 | 0.20 | 0.29 | 0.26 |
| SLC6A8 | 0.30 | 0.22 | 0.26 | 0.26 |
| HELLS | 0.20 | 0.22 | 0.35 | 0.26 |
| C11orf82 | 0.24 | 0.12 | 0.41 | 0.26 |
| S100A9 | 0.25 | 0.29 | 0.22 | 0.26 |
| KDELC2 | 0.31 | 0.17 | 0.28 | 0.26 |
| EME1 | 0.13 | 0.35 | 0.29 | 0.26 |
| TRIM56 | 0.35 | 0.39 | 0.03 | 0.26 |
| VTI1A | 0.46 | 0.03 | 0.27 | 0.26 |
| IER5L | 0.25 | 0.23 | 0.29 | 0.26 |
| CDC25A | 0.22 | 0.25 | 0.30 | 0.26 |
| CDC42BPB | 0.25 | 0.39 | 0.13 | 0.26 |
| TICRR | 0.21 | 0.30 | 0.26 | 0.26 |
| GBP3 | 0.41 | 0.18 | 0.18 | 0.26 |
| MCM5 | 0.34 | 0.27 | 0.16 | 0.26 |
| UBA6 | 0.42 | 0.15 | 0.19 | 0.26 |
| ZNF532 | 0.54 | 0.02 | 0.21 | 0.26 |
| CDC42EP2 | 0.16 | 0.27 | 0.34 | 0.26 |
| KIF5B | 0.25 | 0.25 | 0.27 | 0.26 |
| CHST3 | 0.34 | 0.20 | 0.23 | 0.26 |
| PEX26 | 0.34 | 0.48 | -0.06 | 0.26 |
| S100A2 | 0.37 | 0.24 | 0.15 | 0.25 |
| EMR2 | 0.41 | 0.18 | 0.18 | 0.25 |
| TMED7-TICAM2 | 0.39 | 0.13 | 0.25 | 0.25 |
| SMG5 | 0.12 | 0.42 | 0.22 | 0.25 |
| KIF15 | 0.27 | 0.18 | 0.32 | 0.25 |
| TNFRSF6B | 0.42 | 0.13 | 0.21 | 0.25 |
| TNFRSF9 | 0.48 | 0.09 | 0.19 | 0.25 |

| | | | | |
|---|---|---|---|---|
| WARS | 0.47 | 0.14 | 0.16 | 0.25 |
| HIST2H3A | 0.19 | 0.36 | 0.21 | 0.25 |
| FAM208B | 0.24 | 0.26 | 0.27 | 0.25 |
| ANKRD9 | 0.29 | 0.31 | 0.16 | 0.25 |
| MTCH2 | 0.20 | 0.24 | 0.32 | 0.25 |
| EFHD2 | 0.26 | 0.22 | 0.28 | 0.25 |
| IL1RN | 0.27 | 0.34 | 0.15 | 0.25 |
| MAP2K1 | 0.32 | 0.36 | 0.09 | 0.25 |
| SGOL1 | 0.27 | 0.13 | 0.36 | 0.25 |
| URB2 | 0.28 | 0.23 | 0.25 | 0.25 |
| ELOVL6 | 0.32 | 0.13 | 0.31 | 0.25 |
| CMAS | 0.31 | 0.22 | 0.22 | 0.25 |
| MLF1IP | 0.26 | 0.22 | 0.28 | 0.25 |
| POSTN | 0.50 | 0.06 | 0.20 | 0.25 |
| KLRC2 | 0.32 | 0.15 | 0.28 | 0.25 |
| PXDN | 0.46 | 0.19 | 0.11 | 0.25 |
| AKR1B15 | 0.25 | 0.32 | 0.18 | 0.25 |
| INF2 | 0.23 | 0.36 | 0.16 | 0.25 |
| LINC00346 | 0.37 | 0.12 | 0.26 | 0.25 |
| YWHAG | 0.35 | 0.24 | 0.17 | 0.25 |
| AKR1B10 | 0.26 | 0.33 | 0.16 | 0.25 |
| PSME3 | 0.35 | 0.36 | 0.04 | 0.25 |
| TNFRSF1A | 0.20 | 0.23 | 0.32 | 0.25 |
| PTTG1 | 0.33 | 0.14 | 0.29 | 0.25 |
| DUSP4 | 0.40 | 0.12 | 0.23 | 0.25 |
| DSP | 0.31 | 0.18 | 0.25 | 0.25 |
| FAM114A1 | 0.32 | 0.31 | 0.12 | 0.25 |
| CISD1 | 0.28 | 0.19 | 0.27 | 0.25 |
| JOSD1 | 0.27 | 0.49 | -0.02 | 0.25 |
| TMEM158 | 0.39 | 0.14 | 0.22 | 0.25 |
| WIPI1 | 0.47 | 0.31 | -0.02 | 0.25 |
| GRAMD1B | 0.13 | 0.31 | 0.30 | 0.25 |
| ADRBK1 | 0.18 | 0.24 | 0.33 | 0.25 |
| MAP3K10 | 0.15 | 0.38 | 0.23 | 0.25 |
| ENO2 | 0.35 | 0.10 | 0.29 | 0.25 |
| CDKN2D | 0.34 | 0.25 | 0.16 | 0.25 |
| BORA | 0.25 | 0.27 | 0.23 | 0.25 |
| FAM207A | 0.24 | 0.33 | 0.18 | 0.25 |
| SRGAP1 | 0.23 | 0.26 | 0.25 | 0.25 |
| ADAMTS2 | 0.38 | 0.20 | 0.16 | 0.25 |
| WDR62 | 0.21 | 0.35 | 0.19 | 0.25 |
| PATL1 | 0.22 | 0.13 | 0.40 | 0.25 |
| CENPH | 0.25 | 0.21 | 0.28 | 0.25 |
| HIST2H3C | 0.19 | 0.36 | 0.20 | 0.25 |
| USP5 | 0.18 | 0.34 | 0.22 | 0.25 |

| | | | | |
|---|---|---|---|---|
| SH2D5 | 0.15 | 0.35 | 0.25 | 0.25 |
| ANXA2 | 0.34 | 0.41 | 0.00 | 0.25 |
| ADCY3 | 0.43 | 0.20 | 0.12 | 0.25 |
| PITPNC1 | 0.33 | 0.18 | 0.24 | 0.25 |
| SNX7 | 0.35 | 0.15 | 0.25 | 0.25 |
| FSCN1 | 0.30 | 0.22 | 0.22 | 0.25 |
| UQCRFS1 | 0.27 | 0.50 | -0.03 | 0.25 |
| OAS1 | 0.30 | 0.22 | 0.22 | 0.25 |
| POLA2 | 0.29 | 0.05 | 0.41 | 0.25 |
| TCIRG1 | 0.19 | 0.33 | 0.21 | 0.25 |
| ERRFI1 | 0.17 | 0.26 | 0.32 | 0.25 |
| PARPBP | 0.28 | 0.19 | 0.28 | 0.25 |
| LIMK1 | 0.40 | 0.25 | 0.09 | 0.25 |
| KCNK6 | 0.29 | 0.27 | 0.18 | 0.25 |
| ZNF259 | 0.32 | 0.07 | 0.35 | 0.25 |
| LRR1 | 0.33 | 0.15 | 0.26 | 0.25 |
| RECQL | 0.34 | 0.18 | 0.22 | 0.25 |
| DDX11 | 0.23 | 0.23 | 0.28 | 0.25 |
| ATP5C1 | 0.19 | 0.30 | 0.25 | 0.25 |
| ADAMTS14 | 0.34 | 0.22 | 0.18 | 0.25 |
| CDCA4 | 0.29 | 0.19 | 0.26 | 0.25 |
| MCAM | 0.32 | 0.16 | 0.26 | 0.25 |
| MUC16 | 0.27 | 0.32 | 0.14 | 0.25 |
| FAM83F | 0.20 | 0.38 | 0.15 | 0.25 |
| RNF26 | 0.25 | 0.21 | 0.28 | 0.25 |
| CENPN | 0.21 | 0.26 | 0.27 | 0.25 |
| WDR4 | 0.27 | 0.37 | 0.10 | 0.25 |
| ACOT9 | 0.37 | 0.34 | 0.02 | 0.25 |
| GALNT14 | 0.32 | 0.26 | 0.16 | 0.25 |
| RNF213 | 0.34 | 0.26 | 0.13 | 0.25 |
| PHF19 | 0.32 | 0.23 | 0.19 | 0.25 |
| ESM1 | 0.38 | 0.17 | 0.19 | 0.25 |
| LHFPL2 | 0.31 | 0.27 | 0.15 | 0.25 |
| NAV1 | 0.54 | -0.02 | 0.22 | 0.25 |
| RNASEH1 | 0.32 | 0.27 | 0.14 | 0.25 |
| CARS | 0.29 | 0.17 | 0.27 | 0.25 |
| MND1 | 0.18 | 0.21 | 0.34 | 0.25 |
| GNPNAT1 | 0.22 | 0.16 | 0.35 | 0.25 |

Supplementary **Figure S1**

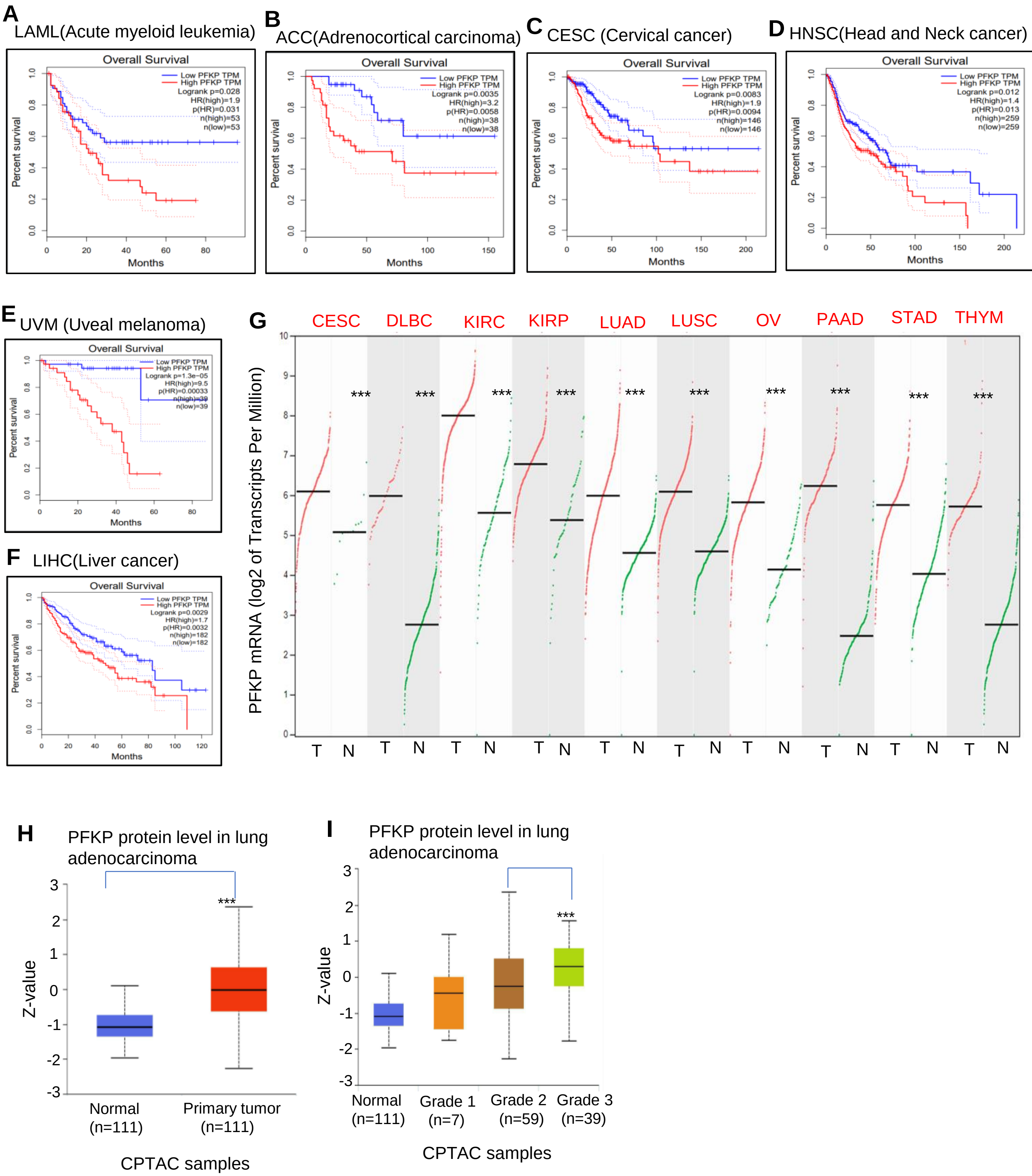

**Figure S1. PFKP is highly expressed in multiple types of cancers and correlates to poor patient survival. A-F**, High levels of PFKP mRNA are related to poor patient survival in multiple types of tumors in TCGA data from the GEPIA website; **G**, PFKP mRNAs are highly expressed in multiple types of tumors in TCGA data from the GEPIA website. CESC, Cervical squamous cell carcinoma and endocervical adenocarcinoma, DLBC, Lymphoid Neoplasm Diffuse Large B-cell Lymphoma, KIRC, Kidney renal clear cell carcinoma, KIRP, Kidney renal papillary cell carcinoma, PAAD, Pancreatic adenocarcinoma; STAD, Stomach adenocarcinoma, THYM, Thymoma; **H, I**, PFKP protein expression was higher in lung adenocarcinomas as compared to normal lung tissues and also higher in poorly differentiated tumors in data from UALCAN website.

Supplementary **Figure S2**

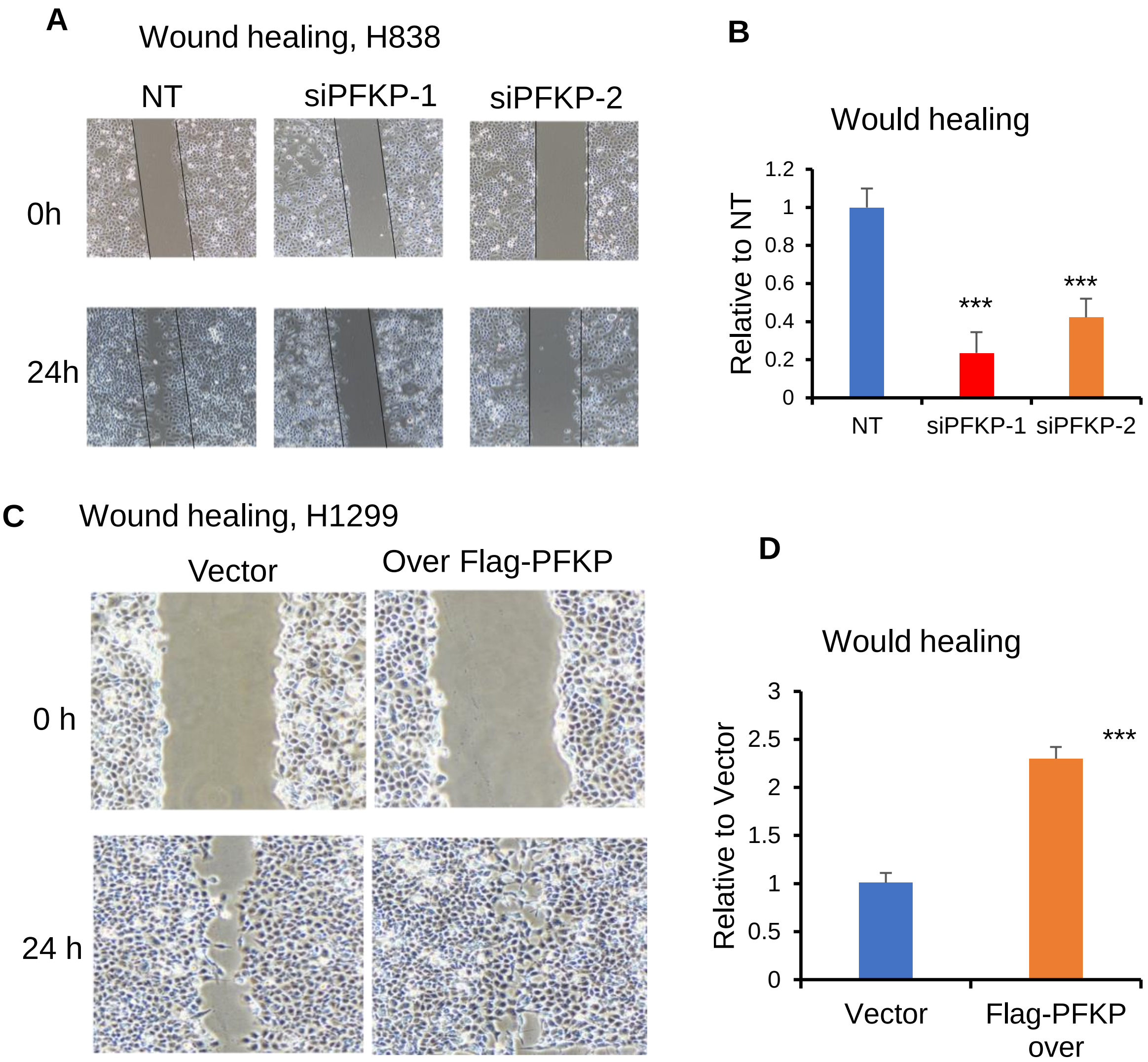


**Figure S2. PFKP affects cell wound healing. A, B,** Silencing of PFKP leads to decreased wound healing in H838 cells, B is the relative value, *** p<0.001; **C, D**, Cells have a higher speed of wound healing upon overexpression of PFKP in H1299 cells as compared to vector.

Supplementary **Figure S3**

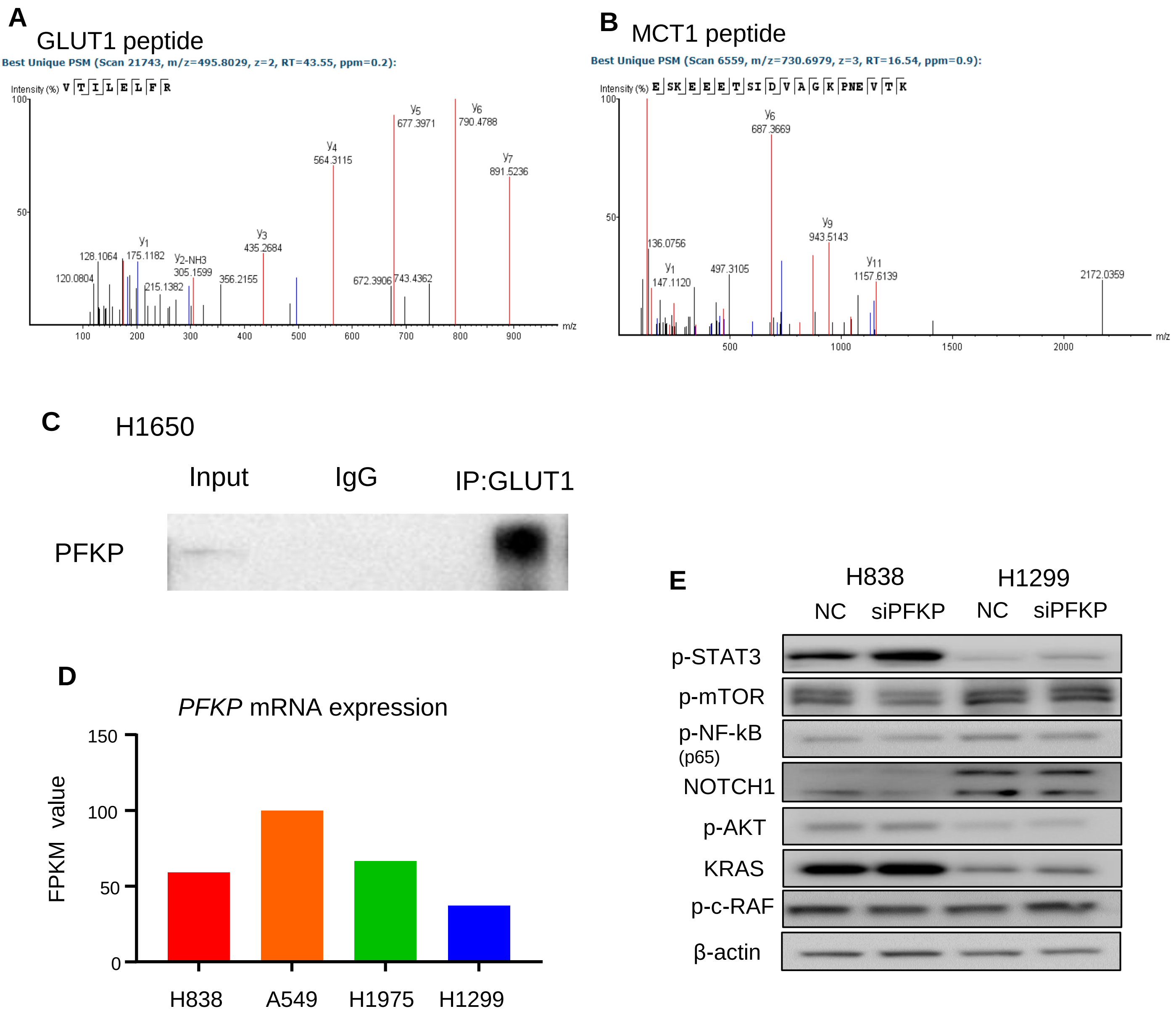


**Figure S3. The images of the mass spectrum from PFKP Co-IP followed by mass spectrometry and Western blotting results**. **A**, The image of GLUT1 peptide; **B**, The image of MCT1 peptide; **C**, Western blot after pull down using anti-GLUT1 antibody Co-IP verified that PFKP binds to GLUT1 directly; **D,** PFKP mRNA level in lung cancer cell lines measured by RNA-seq; **E**, Proteins were not changed upon PFKP knockdown in lung cancer cell lines measured by Western blotting.

Supplementary **Figure S4**

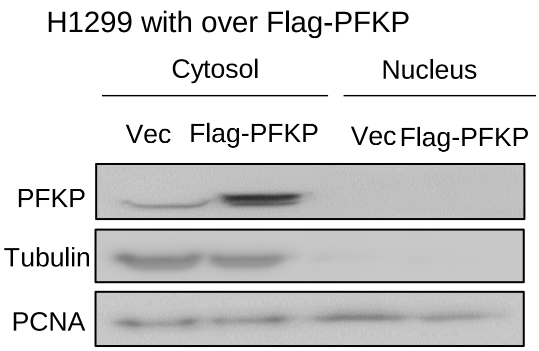


**Figure S4**, PFKP has no expression in the nucleus (H1299 with over Flag-PFKP) measured by Western blotting.

Supplementary **Figure S5**

PFKP TMA 180 IHC 2022 09 09

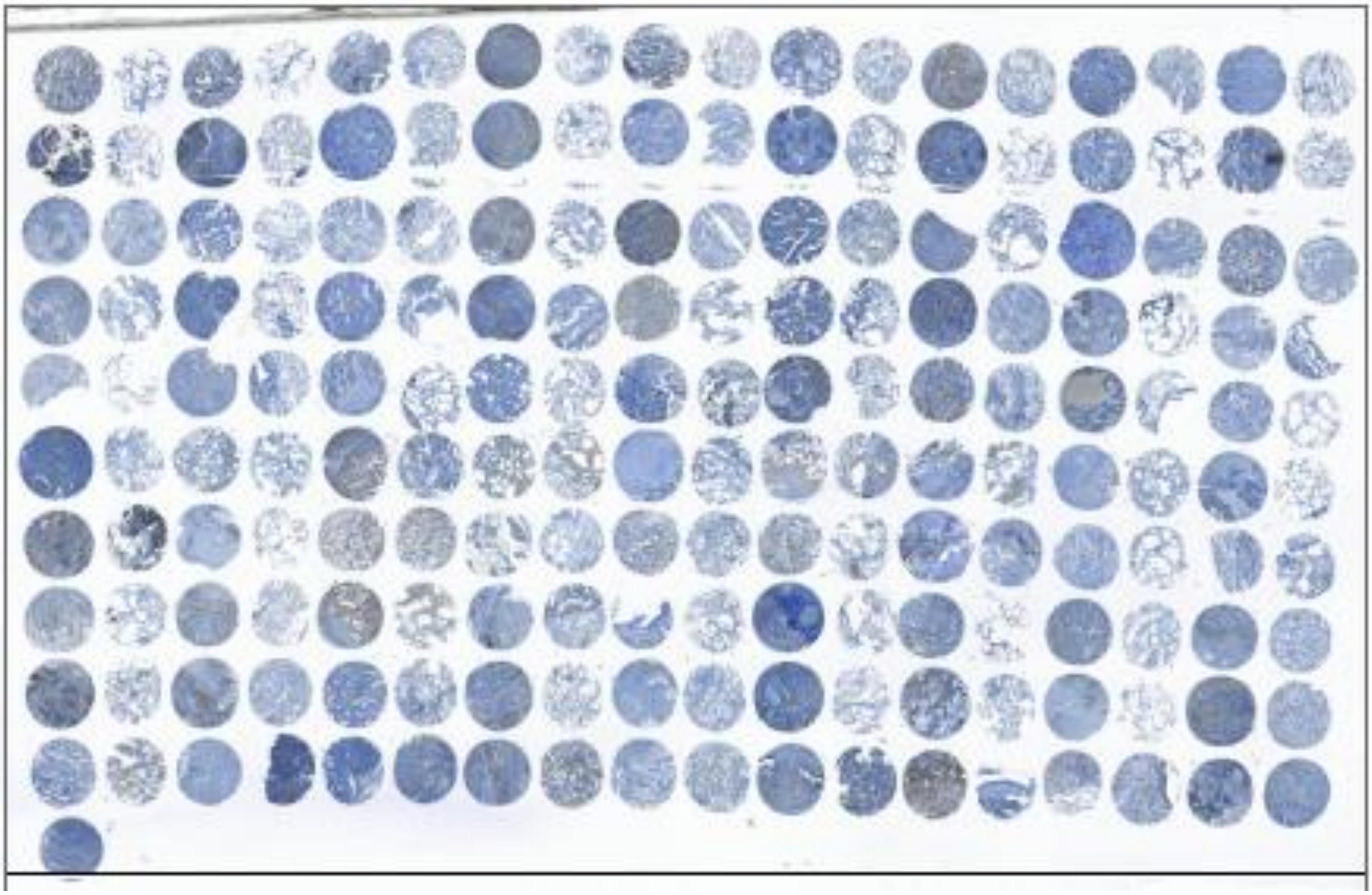

T N T N T N T N T N T N T N T N T N

**Figure S5**, IHC of TMA image from lung adenocarcinoma. T, tumor tissue; N, normal lung tissue

Supplementary **Figure S6**

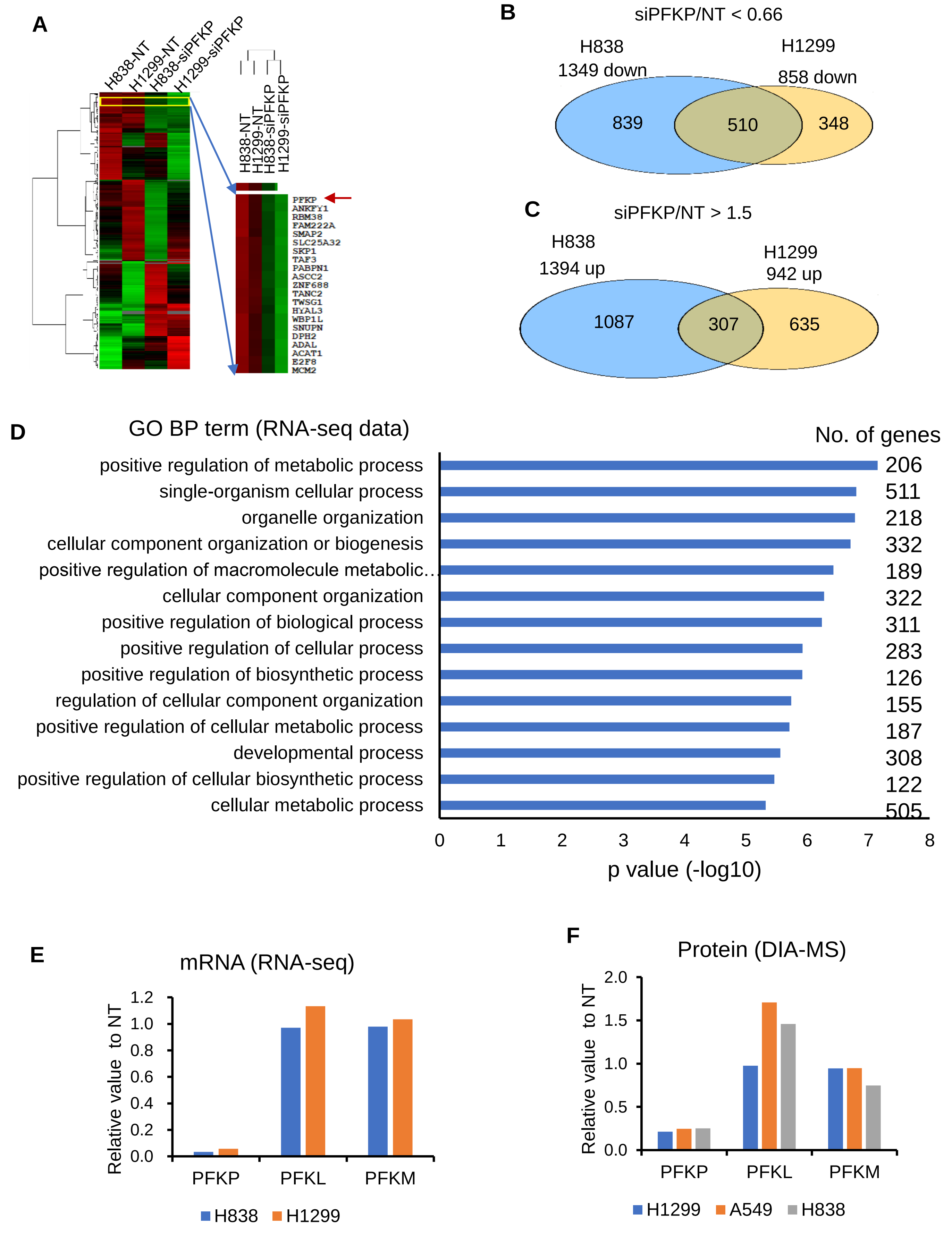

**Figure S6. RNA-seq and proteomics analyses uncover cancer related signaling and metabolic pathways affected by PFKP knockdown.** **A**, Heat map of the changed genes upon PFKP knockdown and control for H838 and H1299 cell lines. **B,** There are 510 down-regulated genes after PFKP knockdown for both H838 and H1299 cell lines. The threshold is fold-change less than 0.66 of siPFKP/control (NT). **C,** There are 307 up-regulated genes after PFKP knockdown for both H838 and H1299 cell lines. The threshold is fold-change larger than 1.5 of siPFKP/control (NT). **D** The 510 down-regulated genes and 307 up-regulated genes were enriched in important metabolism-related signaling pathways; **E** and **F**, Changes of PFKL and PFKM after PFKP knockdown measured by RNA-seq and DIA-MS.

Supplementary **Figure S7**

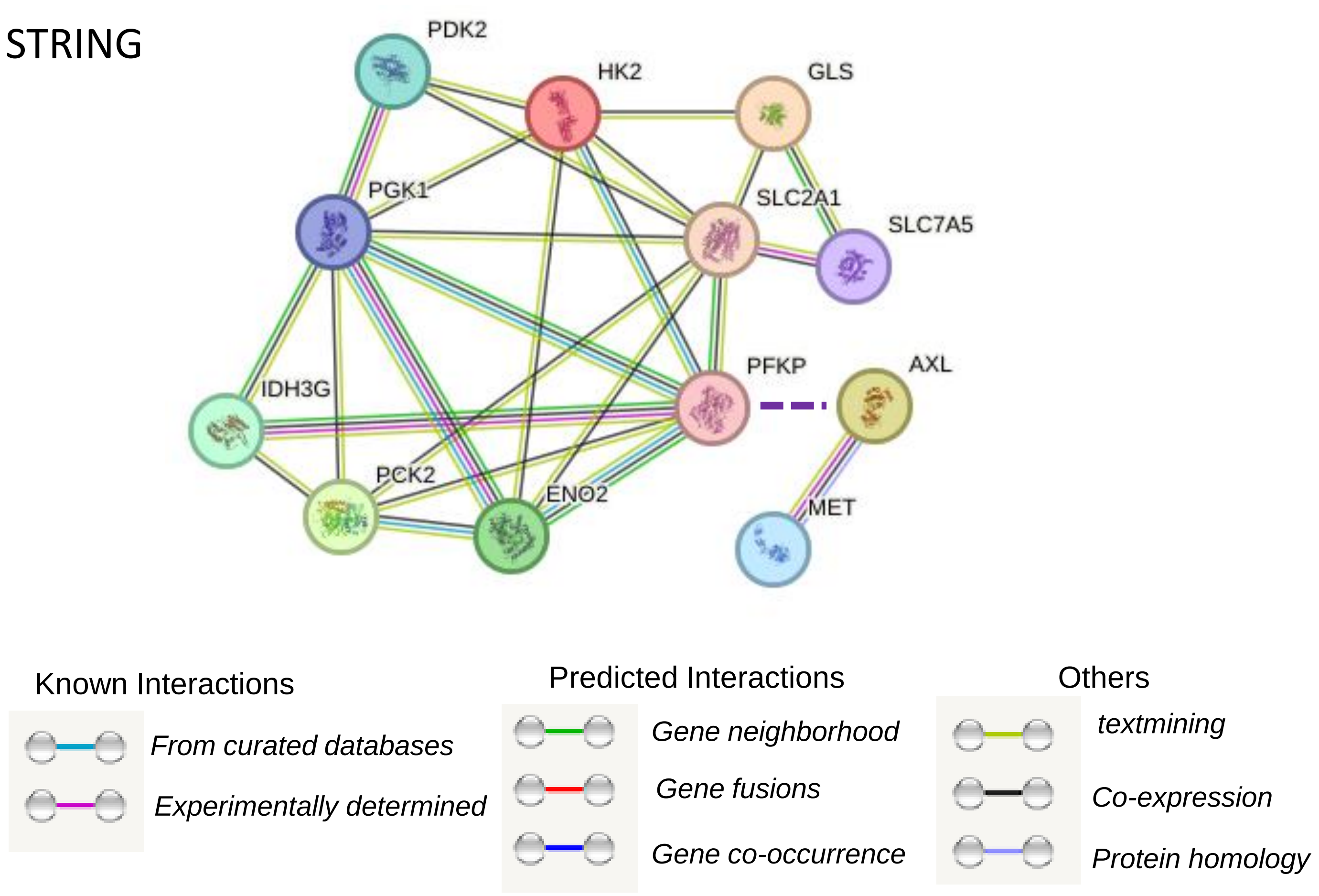


**Figure S7. PFKP-related protein-protein interaction networks.** PFKP-regulated genes/proteins uncovered in this study have potential protein-protein interaction networks using the String website (https://cn.string-db.org/). PFKP may directly interact with AXL.
